# Double mutation of open syntaxin and UNC-18 P334A leads to excitatory-inhibitory imbalance and impairs multiple aspects of *C. elegans* behavior

**DOI:** 10.1101/2023.08.18.553709

**Authors:** Mengjia Huang, Ya Wang, Chun Hin Chow, Karolina P. Stepien, Karen Indrawinata, Junjie Xu, Peter Argiropoulos, Xiaoyu Xie, Kyoko Sugita, Chi-Wei Tien, Soomin Lee, Philippe P. Monnier, Josep Rizo, Shangbang Gao, Shuzo Sugita

## Abstract

SNARE and Sec/Munc18 proteins are essential in synaptic vesicle exocytosis. Open form t-SNARE syntaxin and UNC-18 P334A are well-studied exocytosis-enhancing mutants. Here we investigate the interrelationship between the two mutations by generating double mutants in various genetic backgrounds in *C. elegans*. While each single mutation rescued the motility of *CAPS/unc-31* and *synaptotagmin/snt-1* mutants significantly, double mutations unexpectedly worsened motility or lost their rescuing effects. Electrophysiological analyses revealed that simultaneous mutations of open syntaxin and gain-of-function P334A UNC-18 induces a strong imbalance of excitatory over inhibitory transmission. In liposome fusion assays performed with mammalian proteins, the enhancement of fusion caused by the two mutations individually was abolished when the two mutations were introduced simultaneously, consistent with what we observed in *C. elegans*. We conclude that open syntaxin and P334A UNC-18 do not have additive beneficial effects, and this extends to *C. elegans’* characteristics such as motility, growth, offspring bared, body size, and exocytosis, as well as liposome fusion in vitro. Our results also reveal unexpected differences between the regulation of exocytosis in excitatory versus inhibitory synapses.

## Introduction

Sinusoidal *C. elegans* movement requires coordination between the excitation and inhibition of the neuromuscular junction (Jospin et al., 2009). Excitation/inhibition is initiated by the release of excitatory (acetylcholine) or inhibitory (gamma-aminobutyric acid, GABA) transmitters at their respective synapses via synaptic vesicle exocytosis (Richmond and Jorgensen, 1999).

Synaptic vesicle exocytosis consists of spontaneous and evoked exocytosis. While the fundamental mechanisms of exocytosis are well studied, the mechanisms that are specific to the types of synapses (excitatory vs. inhibitory) or types of exocytosis (spontaneous vs. evoked) are largely unknown. One notable exception is the complexin/CPX-1 protein, which in invertebrates impairs spontaneous exocytosis while enhancing evoked exocytosis (Huntwork and Littleton, 2007; Hobson et al., 2011; Martin et al., 2011).

SNARE (soluble NSF-attachment receptor) and Sec/Munc18 (SM) proteins are the essential factors that govern synaptic vesicle exocytosis. These proteins are conserved and play critical roles regardless of the types of synapses or types of exocytosis. The SNARE proteins syntaxin (UNC-64 in *C. elegans*), synaptobrevin/VAMP and SNAP-25 come together to form a four-helix bundle called the SNARE complex that brings the membranes together, which is key for membrane fusion (Hanson et al., 1997; Sutton et al., 1998; Weber et al., 1998; Rizo, 2022). In this complex, syntaxin and synaptobrevin each contribute one helix to the four-helix bundle, while SNAP-25 contributes two helices (Sutton et al., 1998). The SNARE complex is tightly regulated by SNARE regulatory proteins, such as Munc18-1 (UNC-18 in *C. elegans*) (Hata et al., 1993; Verhage et al., 2000; Weimer et al., 2003), Munc13s (UNC-13 in *C. elegans*) (Brose et al., 1995; Augustin et al., 1999; Richmond et al., 1999) and the Ca^2+^ sensor synaptotagmin-1 (SNT-1 in *C. elegans*) (Geppert et al., 1994; Jorgensen et al., 1995; Li et al., 2021). Together, these proteins work in harmony to fuse vesicles to the plasma membrane and release neurotransmitters in a Ca^2+^-dependent manner (Geppert et al., 1994; Fernández-Chacón et al., 2001).

Loss-of-function mutations of these essential exocytosis proteins lead to death or severe movement defects in *C. elegans,* mice, and humans. On the other hand, gain-of-function mutations lead to enhanced exocytosis and can rescue the phenotypes of other exocytosis defective mutants. The investigation of such gain-of-function mutants provides mechanistic insights into how these proteins and their interactions with other exocytotic proteins contribute to exocytosis. Thus, an L166A/E167A mutation in syntaxin/UNC-64 (abbreviated as “LE”) and a P334A mutation in UNC-18 (P335A mutation in mammalian Munc18-1; abbreviated as “PA”) are well-known gain-of-function mutations that have been extensively studied in various model systems of exocytosis, including *in vitro* liposome fusion (Parisotto et al., 2014; Sitarska et al., 2017), PC12 cells (Han et al., 2014), *C. elegans* (Richmond et al., 2001; Park et al., 2017; Tien et al., 2020) and mice (Gerber et al., 2008; Munch et al., 2016). Both mutants exhibited enhanced fusion/exocytosis in every assay system tested. However, the interaction between the two mutations has not been investigated.

The t-SNARE syntaxin/UNC-64 exists in two conformations, the open and the closed conformations (Dulubova et al., 1999). The LE mutation leaves syntaxin constitutively open, thus facilitating SNARE complex formation (Dulubova et al., 1999; Gerber et al., 2008) and was originally considered to selectively rescue *unc-13* and *unc-10/RIM* mutants in *C. elegans*, without enhancing exocytosis on its own (Richmond et al., 2001). This led to the hypothesis that the role of Munc13/UNC-13 is to open syntaxin at the synapse (Koushika et al., 2001; Richmond et al., 2001), which was later supported by biophysical assays in vitro (Ma et al., 2011). However, later work found that open syntaxin 1B knock-in (KI) mice exhibit enhanced spontaneous and evoked release from cortical neurons (Gerber et al., 2008). Similarly, our recent generation of open syntaxin knock-in worms revealed that this gain-of-function mutation enhances excitatory synaptic transmission on its own and it can rescue a variety of exocytosis-defective mutants, including *synaptotagmin-1/snt-1*, *unc-2* and *CAPS*/*unc-31*, as well as *unc-13 and unc-10* in *C. elegans* (Tien et al., 2020). These findings showed that the facilitation of SNARE complex assembly caused by the LE mutation enhances neurotransmitter release in a variety of genetic backgrounds (Dulubova et al., 1999; Ma et al., 2011).

Munc18/UNC-18 is a multidomain protein (Misura et al., 2000) that plays an important role in the regulation of neurotransmitter release across many species (Hosono et al., 1992; Harrison et al., 1994; Verhage et al., 2000; Voets et al., 2001; Weimer et al., 2003). Munc18/UNC-18 has an arched structure comprised of three domains that form a central cavity. This central cavity is where syntaxin binds when syntaxin is in a closed conformation (Misura et al., 2000). Formation of this binary complex mediates syntaxin trafficking/chaperoning to the plasma membrane (Arunachalam et al., 2008; Han et al., 2009; Han et al., 2011; Han et al., 2014). In addition to syntaxin, Munc18/UNC-18 can bind to synaptobrevin, forming a template for SNARE complex assembly (Parisotto et al., 2014; Baker et al., 2015). The P334A mutation in Munc18/UNC-18 was designed to extend a helix that binds to synaptobrevin and thus enhance binding, but such enhancement was not observed (Parisotto et al., 2014), and synaptobrevin binding does not require extension of this helix (Stepien et al., 2022). Instead, the P334A mutation weakens syntaxin binding (Han et al. 2014) and causes a gain-of-function because the release of contacts between syntaxin and Munc18/UNC-18 at this site is important to form the template complex and initiate SNARE complex assembly (Stepien et al. 2022). Thus, similar to the open syntaxin, P334A knock-in leads to enhanced excitatory synaptic transmission on its own, and rescues *unc-31* and *unc-13* mutant worms (Park et al., 2017).

In the present study, we investigated how the open syntaxin and P334A UNC-18 mutations interact with each other using *C. elegans* as a model system. We initially anticipated that the double mutant would exhibit additive or synergistic effects on exocytosis, as well as enhanced ability to rescue motility and other altered characteristics observed in various *C. elegans* exocytosis mutants. To our surprise, the double mutants exhibit suppressed motility with enhanced acetylcholine release, as measured by thrashing assays and the sensitivity to the acetylcholinesterase inhibitor (aldicarb), respectively. Strikingly, the presence of both gain-of-function mutations within the same worm did not provide benefits to worm size, growth speed, or number of offspring given. Therefore, we further analyzed these single and double mutants by electrophysiology in detail. Our results show that these mutations have differential effects on exocytosis depending on the type of synapse and the type of exocytosis.

## Results

### P334A UNC-18, but not the absence of TOM-1, rescues motility, acetylcholine release, and growth speed of *snt-1* null mutants like open syntaxin

We previously showed that an open syntaxin *unc-64(LE)* KI mutant can rescue the defects in motility and acetylcholine release of various exocytosis mutants, including *snt-1*, *unc-31*, *unc-13* and *unc-10* (Tien et al., 2020). Although we showed that the *unc-18(PA)* KI mutant rescues *unc-31* and *unc-13* mutants (Park et al., 2017), whether *unc-18(PA)* has the ability to rescue a wide-range of exocytosis mutants remains unknown.

Synaptotagmin-1, encoded by the *snt-1* gene in *C. elegans,* functions as a Ca^2+^ sensor for synchronous neurotransmitter release (Geppert et al., 1994). *Snt-1(md290)* null worms show resistance to aldicarb and have smaller body sizes relative to wild-type worms (Jorgensen et al., 1995; Li et al., 2021). Here, we first examined whether *unc-18(PA)* can rescue *snt-1* null worms, like open syntaxin. We also tested whether absence of the inhibitory protein tomosyn (*tom-1* null, *ok285*) can rescue *snt-1* mutants. The TOM-1 protein, encoded by the *tom-1* gene, inhibits spontaneous and evoked release via the formation of an inhibitory tomosyn-SNARE complex (Fujita et al., 1998). Moreover, the *tom-1(ok285)* null mutation is known to enhance acetylcholine release and partially rescues *unc-13* mutants (McEwen et al., 2006).

Knock-in *C. elegans* strains bearing either the open syntaxin *unc-64(LE)* or *unc-18(PA)* mutations were generated as previously reported (Park et al., 2017; Tien et al., 2020). These mutant worms, along with *tom-1(ok285)* null worms, were subsequently crossed with synaptotagmin null worms, s*nt-1(md290)*. Worms were assayed for their motility and acetylcholine release ability through thrashing and aldicarb sensitivity assays, respectively. We found that both *snt-1; unc-64(LE)* and *snt-1; unc-18(PA)* double mutants similarly and significantly increased thrashing compared to the *snt-1* single mutant (Fig. 1A). Conversely, *tom-1; snt-1* double mutants did not significantly improve the motility of *snt-1* null worms (Fig. 1A). Aldicarb sensitivity assays revealed that *snt-1(md290)* null worms exhibit minimal paralysis after 4 hours (Fig. 1B), suggesting a strongly impaired acetylcholine release. On the other hand, as previously shown, *snt-1*; *unc-64(LE)* double mutants showed increased aldicarb sensitivity compared with *snt-1*, suggesting an increase in acetylcholine release (Tien et al., 2020). Moreover, we found a similar increase of acetylcholine release by the *snt-1; unc-18(PA)* double mutant (Fig. 1B). However, only a slight rescue was seen by the *snt-1; tom-1* null mutant (Fig. 1C).

**Figure 1.**
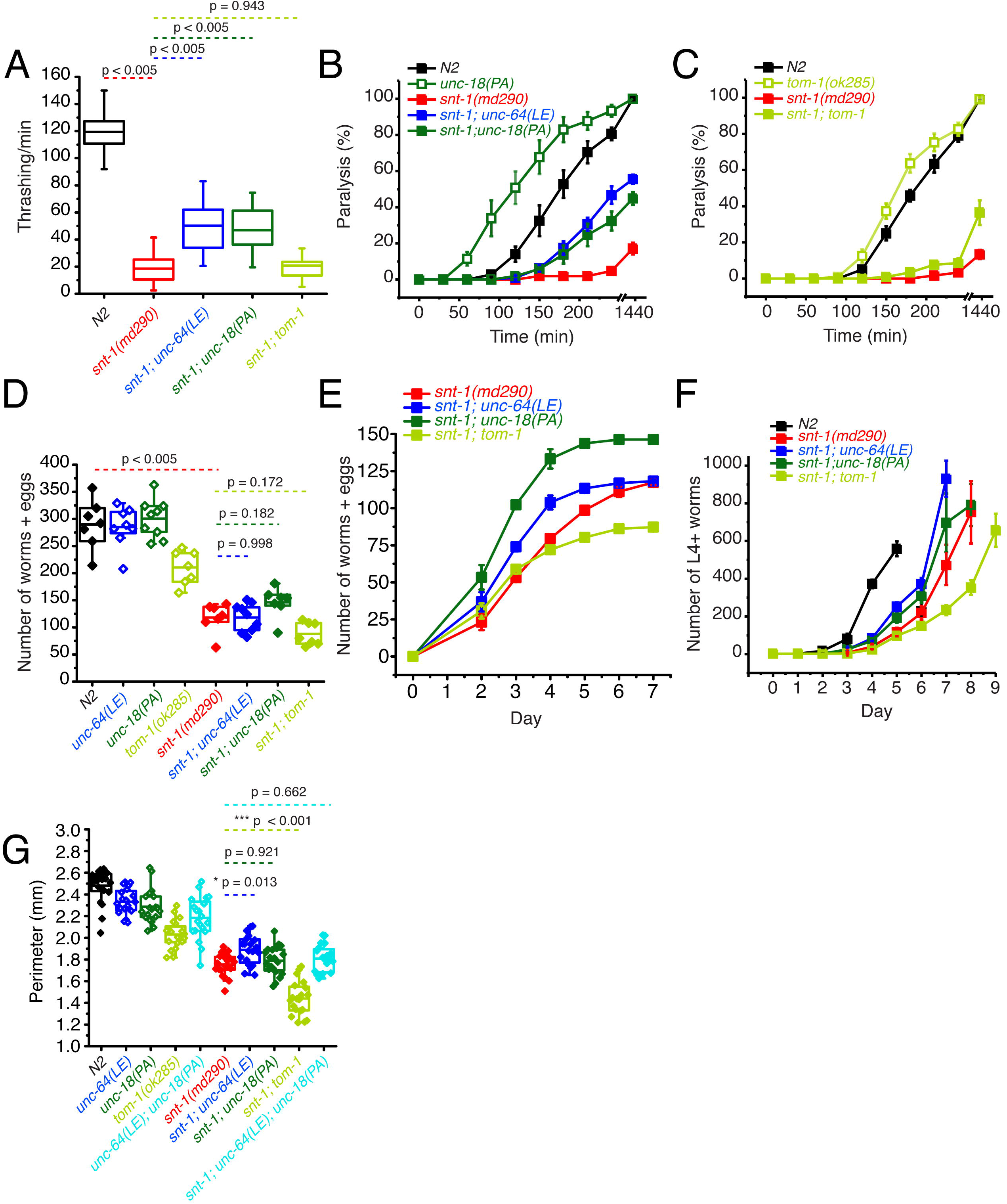
P334A UNC-18, open syntaxin, but not the absence of *tom-1*, rescues many deficits observed in the *snt-1* null mutant. (A) Box and whisker plot of thrashing for N2 control (black), *snt-1* null (red), and *snt-1* double mutants (with *unc-64(LE)*, blue; *unc-18(PA)*, dark green; *tom-1(ok285)*, light green). One-way ANOVA was performed (F_(4, 195)_ = 359, p = 0.00). *snt-1* null worms show decreased thrashing (19 thrashes/min) and are significantly rescued by *unc-64(LE)* (50 thrashes/min, Tukey’s test, p < 0.005) and *unc-18(PA)* (47 thrashes/min, p < 0.005), but are not rescued by absence of *tom-1* (21 thrashes/min, p = 0.942). (B) Aldicarb assay of the indicated strains. *snt-1* null (red) shows strong impairment of aldicarb release when compared with N2 (black). The *unc-18(PA)* single mutant (hollow dark green squares) increases aldicarb sensitivity further than N2 levels. Both *snt-1;unc-64(LE)* and *snt-1;unc-18(PA),* blue and solid dark green respectively, rescued the lost aldicarb sensitivity observed in the *snt-1* null worm. (C) Aldicarb assay of snt-1 rescued by absence of *tom-1*. *tom-1* worms (hollow, light green) exhibit increased aldicarb sensitivity. Only a slight rescue is observed in the *snt-1;tom-1* mutant (filled, light green). (E) Brood size of the indicated worm strains, *snt-1* worms (red) decrease in brood size when compared to N2 (black), *snt-1; unc-64(LE) (*in blue) and *snt-1;unc-18(PA)* (in dark green) have a trend for rescuing the brood size of the worm. (F) Number of worms and eggs laid by each worm strain by day, both *snt-1; unc-64(LE)* in blue and *snt-1;unc-18(PA)* in dark green, increase the worms and eggs laid while *snt-1;tom-1* decreases the worms and eggs laid. (G) Population grown of indicated worm strains. N2 worms (black) sharply increased in population size between days 3 and 4 while snt-1 null worms (red) exhibited continuous growth, peaking between days 7 - 9. Both *snt-1;unc-64(LE) and snt-1;unc-18(PA),* blue and dark green respectively, grew faster than snt-1 null worms while *snt-1;tom-1* (light green) grew even slower than snt-1 null worms. (G) Box and whisker plot of worm perimeter data from indicated strains. *snt-1* null worms (red) show a decrease in size, which was rescued by *snt-1;unc-64(LE)* (blue). *snt-1;unc-18(PA) (*in dark green) and snt-1;*unc-64(LE); unc-18(PA)* (in cyan) did not increase the size of the worm. *tom-1 null* worms (light green) exhibited a smaller body size, and again, *snt-1;tom-1* (light green) further decreased the size of the worm.

*Snt-1(md290)* worms have smaller brood sizes, body sizes and exhibit slow population growth speeds (Jorgensen et al., 1995; Li et al., 2021). Therefore, we also tested whether *unc-64(LE)*, *unc-18(PA)*, or *tom-1(ok285)* mutations can rescue the reduced brood size, population growth speed, and body sizes observed in *snt-1* null worms. For brood size, a single L4 worm was placed on an agar plate and after 48 hours of growth, the original worm was transferred to a fresh agar plate. Afterwards, the original worm was transferred to a fresh new plate every 24 hours until the worm died, or egg laying ceased. Offspring (eggs laid and already hatched worms) were counted after the original worm was removed from the plate. We found that *unc-64(LE)* and *unc-18(PA)* worms both had comparable brood sizes to that of N2, while *tom-1* had a significantly reduced brood size (Fig. 1D). *snt-1* null worm brood sizes were even smaller than the brood size of *tom-1* worms (Fig. 1D). When we crossed the various mutants into the *snt-1* null background, although there was a trend for rescue by the *snt-1; unc-18(PA)* double mutant, the double mutation of *snt-1; tom-1* seemed to worsen the small brood size of *snt-1* null worms (Fig. 1D).

For the population growth assay, three worms of each strain were placed on agar plates and allowed to grow over the course of 9 days, and L4 and above adult worms were counted daily. N2 worms showed a sharp increase in total worm number between days 3 and 4, reaching a peak at day 5 (Fig. 1F). *Snt-1* null worms lagged and did not peak in population until day 8 showing a slow rate of population growth. Both *snt-1; unc-64 (LE)* and *snt-1; unc-18(PA)* worm populations grew faster than *snt-1* null worms. The rescue observed by open syntaxin was stronger than that of the P334A UNC-18 mutant; *snt-1; unc-64(LE)* worms reached their peak a day earlier than *snt-1; unc-18(PA)* worms. However, *snt-1; tom-1* worms displayed decreased growth speeds compared to the *snt-1 null* mutant.

Despite finding no significant rescue in brood size in either *snt-1; unc-64(LE)* or *snt-1; unc-18(PA)*, we did observe an increase in populational growth of these two strains when compared to *snt-1* null worms (Fig 1F). This discrepancy could be explained by *snt-1; unc-64(LE)* worms giving birth to a larger portion of their offspring earlier on. Indeed, when we plotted the brood counts by day (Fig. 1E), we observed that, while *snt-1; unc-64(LE)* worms and *snt-1;unc-18(PA)* worms had a similar cumulative brood counts, *snt-1; unc-64(LE)* worms had a left-shifted curve and laid more eggs in the beginning. These worms that were laid earlier (F1 progeny) would then mature and give rise to the F2 progeny that may be reflected in the rapid peak observed in the population growth assay (Fig. 1F).

Next, we analyzed worm body size. For this purpose, *C. elegans* were synchronized and imaged two days after they reached the L4 stage. The captured image of the worm was manually traced to find the perimeter of the worm. We found that *snt-1* null worms showed a smaller body size compared to N2 worms (Fig. 1G). While *unc-64(LE)* weakly increased the size of *snt-1 null* worms, *unc-18(PA)* did not. We found that *tom-1 null* single mutants also exhibited a smaller body size, which is consistent with previous literature (Lee et al., 2011), and *snt-1;tom-1* worms resulted in a further significant decrease to the size of the worm. These results may suggest that hypersecreting mutants such as *tom-1* (Lee et al., 2011), open syntaxin, and P334A UNC-18 (Park et al., 2017) may have a common phenotype of having a smaller body size. Overall, we found that open syntaxin and P334A UNC-18 have similar rescuing abilities on motility, acetylcholine release, brood size and growth speed on *snt-1* null mutants. Such rescuing effects are absent in the *tom-1* null mutant. The results also suggest that open syntaxin and P334A UNC-18 utilize similar mechanisms to enhance exocytosis, which is distinct from the exocytosis-enhancing mechanism exhibited by the lack of the inhibitory tomosyn/TOM-1 protein.

### Simultaneous P334A UNC-18 and open syntaxin mutation abolishes the ability to rescue motility and growth speed of *snt-1* mutants but rescues aldicarb sensitivity

We found that open syntaxin and P334A UNC-18 mutants can individually rescue the motility, brood size, and growth speed of *snt-1* mutants, and that this rescue is accompanied by an increase in acetylcholine release. What remains unknown is if/how the rescuing effects of the two mutants are interrelated. If open syntaxin and P334A UNC-18 use independent mechanisms to enhance exocytosis of *snt-1* null, we would anticipate that the *unc-64(LE) and unc-18(PA)* double mutation may have additive or synergistic effects when rescuing the *snt-1* null mutant. On the other hand, if they use common or similar mechanisms, we may see saturating effects between the two. Therefore, we next asked if the double mutation would exhibit an additional ability to rescue motility and acetylcholine release in the *snt-1* null background. As such, we generated the *snt-1; unc-64(LE); unc-18(PA)* triple mutant. Despite both open syntaxin and P334A UNC-18 mutants being able to rescue thrashing activity individually (Fig. 1A), the *snt-1; unc-64(LE); unc-18(PA)* triple mutant worms showed a similar thrashing count to the *snt-1* null mutant (Fig. 2A). The increased brood size and population growth rate seen by *snt-1; unc-64(LE)* and *snt-1; unc-18(PA)* double mutants were also abolished in the triple mutant – *snt-1; unc-64(LE); unc-18(PA)* worms, which showed a trend for reduced brood size (Fig. 2C), and a similar growth rate to that of the *snt-1* null worm (Fig. 2D). Importantly, however, the triple mutant was still able to show a strong rescue of aldicarb sensitivity (Fig. 2B). Overall, we surprisingly find that the behavioral benefits gained by the single open syntaxin or single P334A UNC-18 mutants are lost when both mutations are present simultaneously; yet they strongly rescued aldicarb sensitivity. Thus, the triple mutant exhibits a striking dissociation between behavior and aldicarb sensitivity.

**Figure 2.**
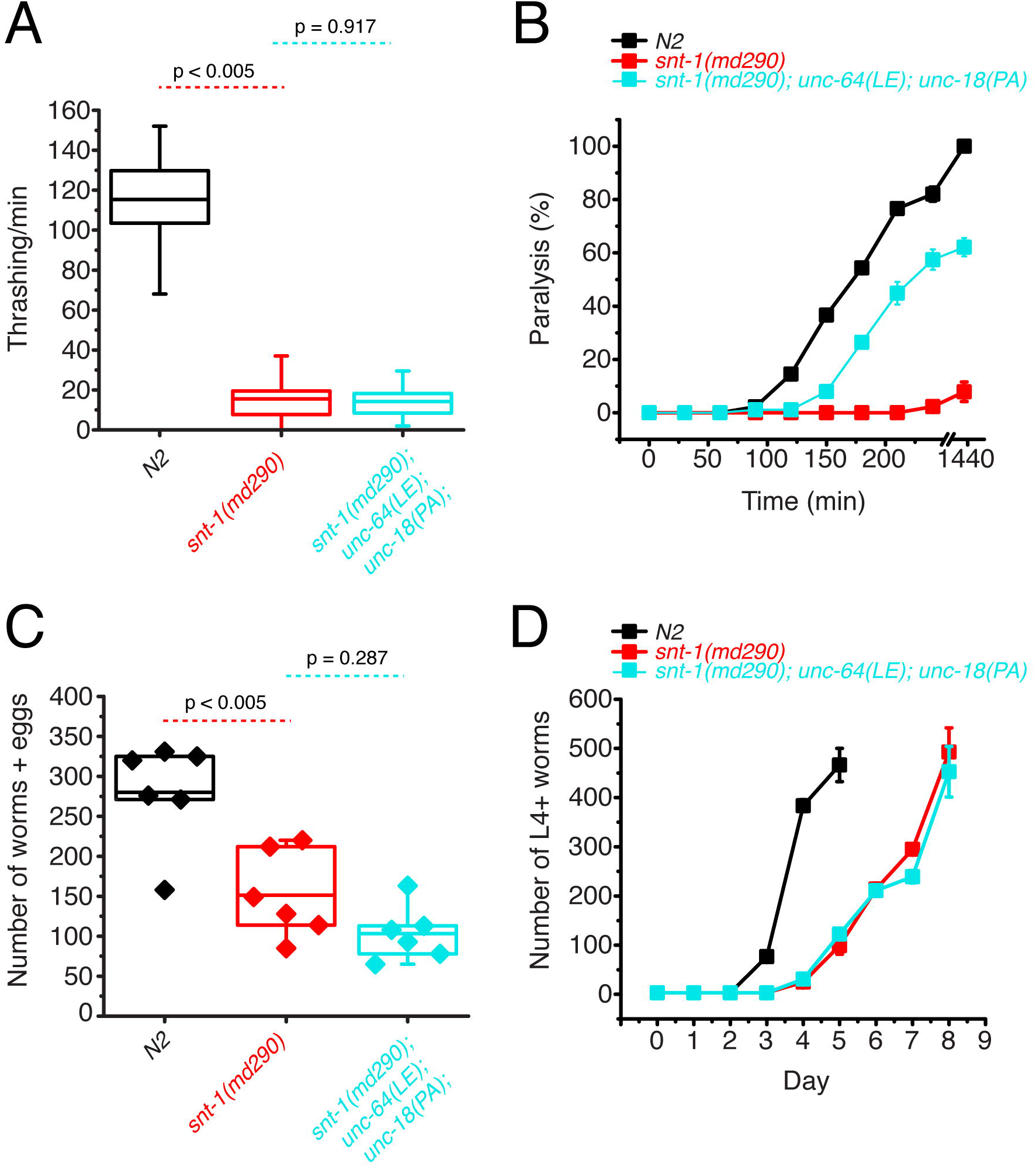
Double mutations of P334A UNC-18 and open syntaxin abolish the ability to rescue motility and growth speed of *snt-1* while they rescue aldicarb sensitivity. (A) Box and whisker plot of thrashing for *snt-1* null (red), *snt-1; unc-64(LE); unc-18(PA)* triple mutant (cyan), and N2 control (black). One-way ANOVA was performed (F_(2, 117)_ = 599, p = 0.00). *snt-1* null worms show decreased thrashing (16 thrashes/min) and are not rescued by the triple mutant (14 thrashes/min, Tukey’s test, p < 0.917). (B) Aldicarb assay of *snt-1* (red), *snt-1; unc-64(LE); unc-18(PA)* triple mutant (cyan), and N2 control (black). A rescue is seen in aldicarb sensitivity by the *snt-1; unc-64(LE); unc-18(PA)* triple mutant worm. (C) Brood size of indicated strains, *snt-1* null worms (red) exhibited a decreased brood size when compared to N2 (black), one-way ANOVA (F_(2, 15)_ = 17.987, p = 0.0001), and the *snt-1; unc-64(LE); unc-18(PA)* triple mutant worms (cyan) trended to decrease (n.s., p = 0.287). (D) Population growth of the indicated worm strains. N2 worms (black) sharply increased in population size between days 3 and 4 while snt-1 null worms (red) exhibited continuous growth, peaking at day 8. *snt-1;unc-64(LE)*;*snt-1;unc-18(PA)* triple mutant worms (cyan) showed a similar population growth trend to that of the *snt-1* null worm.

### The P334A UNC-18 and open syntaxin double mutant dramatically worsens the motility of *unc-31* while rescuing aldicarb sensitivity

We next tested the two knock-in mutations in a different background to see if the loss of beneficial effects of the combined mutations is specific to the *snt-1* null background. CAPS1/UNC-31 has been shown to play key roles in dense core vesicle docking (Hammarlund et al., 2008). In addition, a few studies suggest a significant role for CAPS1/UNC-31 in synaptic vesicle exocytosis (Renden et al., 2001; Jockusch et al., 2007). *Unc-31(e928)* null worms are slow and sluggish, exhibiting a low thrashing rate and impaired aldicarb sensitivity (Charlie et al., 2006). We previously showed that the low thrashing of *unc-31(e928)* was significantly rescued by open syntaxin and P334A UNC-18 (Park et al., 2017; Tien et al., 2020). We originally anticipated that a double mutation of open syntaxin and P334A UNC-18 would further improve the thrashing activity of the *unc-31* null mutant. Contrary to our expectations, simultaneous mutations of *unc-64(LE)* and *unc-18(PA)* in the *unc-31(e928)* background, namely *unc-31(e928); unc-64(LE); unc-18(PA)* triple mutant, strikingly abrogated the motility of the *unc-31* worm (Fig. 3A). Moreover, *unc-31(e928)* worms exhibited a comparable body size to that of N2 worms and both *unc-31; unc-64(LE)* and *unc-31(e928); unc-18(PA)* double mutants slightly increased the body size of the worm (Fig. 3C). However, the *unc-31(e928); unc-64(LE); unc-18(PA)* triple mutant exhibited a significantly smaller body size when compared to the single and either of the double mutant worms. We once again find that the benefits gained by the *unc-64(LE)* and *unc-18(PA)* mutants are lost or even worsened when both mutations are simultaneously present.

**Figure 3.**
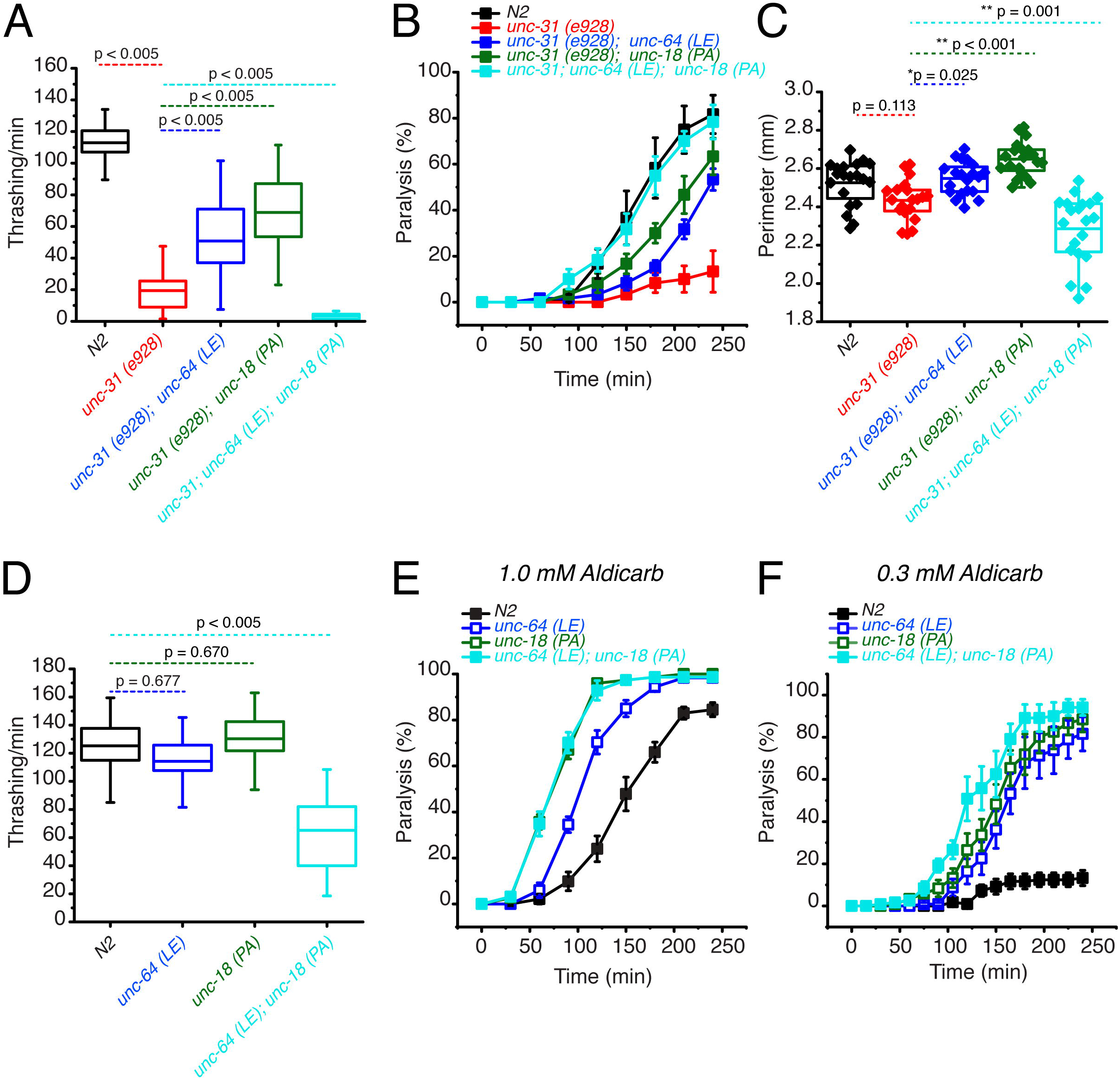
Open syntaxin and P334A unc-18 double mutation combination dramatically worsens the motility of *unc-31* while rescuing aldicarb sensitivity. (A) Box and whisker plot of thrashing for *unc-31* null (red), *unc-31; unc-64(LE)* double mutant (blue)*, unc-31;unc-18(PA)* double mutant (dark green), and *unc-31; unc-64(LE), unc18(PA)* triple mutant (cyan), and N2 control (black). One-way ANOVA was performed (F_(4, 225)_ = 251, p = 0.00). *unc-31* null worms show decreased thrashing (14 thrashes/min) and rescued by both open syntaxin (*unc-31; unc-64(LE),* 51 thrashes/min, Tukey’s test: p < 0.005, blue), P334A unc-18 (*unc-31;unc-18(PA),* 69 thrashes/min, p < 0.005, dark green) but are not rescued by the triple mutant (cyan, 3 thrashes/min, p < 0.005). (B) Aldicarb assay of *unc-31* null (red), *unc-31; unc-64(LE)* double mutant (blue)*, unc-31;unc-18(PA)* double mutant (dark green), and *unc-31; unc-64(LE), unc18(PA)* triple mutant (cyan), and N2 control (black). *unc-31* null worms have a decreased aldicarb sensitivity that is rescued by all mutants tested. The triple mutant rescued aldicarb sensitivity to levels comparable to N2 control. (C) Box and whisker plot of worm perimeter data from indicated strains. unc-31 null (red), *unc-31; unc-64(LE)* double mutant (blue)*, unc-31;unc-18(PA)* double mutant (dark green), and *unc-31; unc-64(LE), unc18(PA)* triple mutant (cyan), and N2 control (black). One-way ANOVA was performed (F_(4, 95)_ = 26.4; p = 0.000): *unc-31* null worms decrease in perimeter length and is rescued by the *unc-31; unc-64(LE)* double mutant (blue), and *unc-31;unc-18(PA)* double mutant (dark green). The *unc-31; unc-64(LE), unc18(PA)* triple mutant (cyan) significantly decreased the perimeter size of the worm. (D) Box and whisker plot of thrashing for *unc-64(LE)* in blue, *unc-18(PA)* in dark green, *unc-64(LE); unc-18(PA)* double mutant in cyan, and N2 in black. One-way ANOVA was performed (F_(3, 156)_ = 90.6, p = 0.000), the *unc-64(LE); unc-18(PA)* double mutant significantly decreased in thrashes per min. (E) Aldicarb assay in 1.0 mM aldicarb of indicated strains, the *unc-64(LE); unc-18(PA)* double mutant had increased aldicarb sensitivity comparable to that of *unc-18(PA)*. (F) Aldicarb assay in 0.3 mM aldicarb of indicated strains, at lower aldicarb concentrations, the *unc-64(LE); unc-18(PA)* double mutant (cyan) showed increased aldicarb sensitivity further than that of *unc-18(PA)* single mutant (dark green).

The loss of aldicarb sensitivity observed in *unc-31(e928)* worms was partially rescued by the *unc-64(LE)* or the *unc-18(PA)* mutants (Fig. 3B). The triple mutant with open syntaxin and P334A UNC-18 increased aldicarb sensitivity even further, exhibiting aldicarb sensitivity comparable to wild-type N2 levels (Fig. 3B). Thus, we observed a striking dissociation between motility and acetylcholine release in the *unc-31(e928); unc-64(LE); unc-18(PA)* triple mutant. This trend is similar to that observed for the *snt*-*1; unc-64(LE); unc-18(PA)* triple mutant.

Since we found detrimental effects in motility when simultaneously expressing *unc-64(LE)* and *unc-18(PA)* mutations in *snt-1* null and *unc-31* null backgrounds, we hypothesized that the double mutation may induce motility defects in a wild-type background. Thus, we crossed *unc-64(LE)* worms with *unc-18(PA)* worms to look at the resulting phenotype of the double mutants in a wild-type background. We observed that both single *unc-64(LE)* and *unc-18 (PA)* mutants thrashed at a degree comparable to N2. However, the *unc-64(LE); unc-18(PA)* double mutants exhibited a dramatically decreased thrashing, with only 65 thrashes/min, which is about half of the wild-type thrashing (Fig. 3D). As previously shown, we found that *unc-64(LE)* or *unc-18(PA)* single mutants display increased aldicarb sensitivity compared to N2 (Park et al., 2017; Tien et al., 2020). At the usual 1 mM concentration of aldicarb, there was no further increase observed by the double mutant of *unc-64 (LE); unc-18(PA)* compared to the *unc-18(PA)* single mutant (Fig. 3E). However, at a lower (0.3 mM) concentration of aldicarb, we observed an increase in aldicarb sensitivity in the double mutant compared to the two single mutants (Fig. 3F). From these experiments (Fig. 1, 2, 3), we conclude that independently, *unc-64(LE)* and *unc-18(PA)* have beneficial effects on the motility, exocytosis, brood, and growth speed of exocytosis-defective mutants, such as *snt-1* or *unc-31* null mutants. However, when these mutants are present simultaneously, their beneficial effects are lost or the mutations become detrimental regardless of the presence or absence of exocytosis-defective mutations. Importantly however, the double mutant increased aldicarb sensitivity irrespective of the background.

### P334A UNC-18 enhances excitatory synaptic transmission like open syntaxin but the enhancement is lost in the double mutant

Why do the double mutants exhibit reduced motility while it increases or rescues aldicarb sensitivity (Fig. 1 - 3)? We envisioned two scenarios for these findings: in the first scenario, we hypothesized that the double mutation causes a severe imbalance of excitation over inhibition; that is, the double mutation selectively enhances acetylcholine release at the excitatory synapse while decreasing GABA release at the inhibitory synapse. In the second scenario, we hypothesized that the double mutation selectively increases spontaneous release while decreasing evoked transmitter release. Such phenomenon was previously shown in *cpx-1 null* mutants, which exhibit reduced motility but enhanced aldicarb sensitivity (Hobson et al., 2011; Martin et al., 2011). Therefore, to reveal the underlying mechanism of the paradoxical phenotype of the double mutant, we decided to examine spontaneous and evoked neurotransmitter release from both excitatory and inhibitory synapses from wild-type control *C. elegans*, the single mutants *unc-64(LE)* and *unc-18(PA)*, and the double *unc-64(LE)*; *unc-18(PA)* mutant. To measure evoked release from either excitatory or inhibitory synapses, we generated the respective mutant lines in the *zxIs6* or *zxIs3* background (Liewald et al., 2008; Tien et al., 2020). *zxIs6* and *zxIs3* strains express channelrhodopsin-2 in excitatory cholinergic neurons (*zxIs6*) or inhibitory GABAergic neurons (*zxIs3*), enabling optogenetic stimulation for accessing transmitter release.

First, recordings were performed in the absence of light stimulation to identify excitatory spontaneous release using the single and double mutant strains generated in a *zxIs6* background. Both *unc-64(LE)* and *unc-18(PA)* single mutants increased the frequency of spontaneous miniature excitatory postsynaptic currents (mEPSCs) (Fig. 4A, C), congruent with what we found previously (Park et al., 2017; Tien et al., 2020). While the *unc-64(LE); unc-18(PA)* double mutant exhibited a trend for increase in mEPSC frequency (Fig. 4A, B, table 1), the averaged value was lower than both of the single mutants and its comparison with WT did not reach statistical significance (p = 0.138). This observation indicates that the *unc-64(LE)* and *unc-18(PA*) mutations do not have additive effects on mEPSC frequency. The amplitude of the spontaneous mEPSCs was not significantly changed across all strains (Fig. 4B). Next, we applied optogenetic stimulation (10 ms, blue light) and measured excitatory evoked postsynaptic currents (EPSCs). Similar to previous reports (Tien et al., 2020), *unc-64(LE)* mutants showed an increase in charge transfer of excitatory postsynaptic currents (EPSCs) without affecting the EPSC amplitude when compared to wild-type worms (Fig. 4D-H). *unc-18(PA)* single mutants exhibited a similar EPSC phenotype to *unc-64(LE)* and increased charge transfer without largely affecting EPSC amplitude (Fig. 4D-H). However, when *unc-64(LE)* and *unc-18(PA)* were present together, the two gain-of-function mutations again seemed to cancel each other. The *unc-64(LE); unc-18(PA)* double mutant only showed a trend for increase in EPSC charge transfer, again yielding values that were lower than those of the single mutants and did not reach statistical significance compared to the WT values (Fig. 4D-H). Thus, the double mutant exhibits a weaker increase in evoked release than either of the single mutants at the excitatory synapse, and also a tendency to lower spontaneous release than the single mutants.

**Figure 4.**
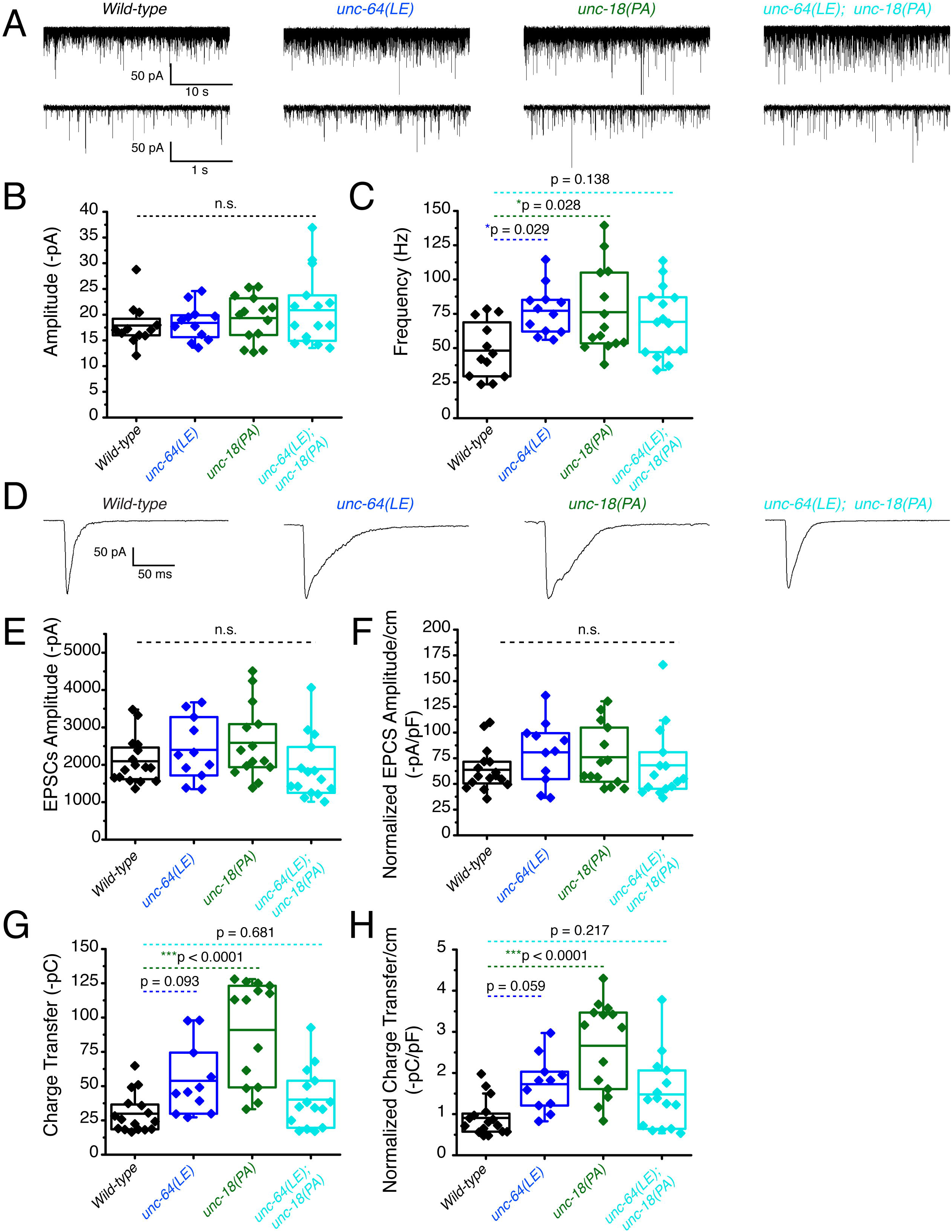
Optogenetic stimulation of P334A UNC-18 and open syntaxin show the double mutant does not facilitate excitatory synaptic transmission like the single mutants. (A) Sample traces of spontaneous miniature postsynaptic currents (mPSCs) of wild-type (WT), open syntaxin (blue), P334A unc-18 (green), and the resulting double mutant (cyan). (B) Amplitude of mPSCs was unchanged in all strains (one-way ANOVA, F_(3, 49)_ = 0.9202; p = 0.4381). (C) Frequency of mPSCs. Both open syntaxin and P334A UNC-18 increased the frequency of mPSCs (one-way ANOVA, F_(3, 49)_ = 3.678; p = 0.0182; open syntaxin: p = 0.0287; P334A UNC-18: p = 0.0281), while a trend of increase was observed in the *unc-64(LE);unc-18(PA)* double mutant, values did not reach significancy (p = 0.1378). (D) Sample traces of evoked excitatory postsynaptic currents (EPSCs) for the indicated strains. (E) EPSC amplitude was unchanged for all strains (one-way ANOVA, F_(3, 52)_ = 2.038; p = 0.1199). (F) EPSC amplitude when calibrated with membrane capacitance to account for differences in animal size also did not show significant changes (one-way ANOVA, F_(3, 52)_ = 0.9039 p = 0.4456). (G) Charge transfer of EPSCs. Both unc-64(LE) and unc-18(PA) single mutants showed an increase in charge transfer (one-way ANOVA, F_(3, 52)_ = 15.76; p <0.0001); however, the unc-64(PA);unc-18(LE) double mutant did not show a significant increase in charge transfer (p = 0.6813). (H) Charge transfer of EPSCs calibrated to membrane capacitance to account for differences in animal size. Charge transfer again is increased for both unc-64(LE) and unc-18(PA) single mutants(F_(3, 52)_ = 11.98; p < 0.0001). A trend of increase is also seen in the EPSC charge transfer in the unc-64(LE);unc-18(PA) double mutant but again did not reach statistical significance (p = 0.2172)

**Table 1:**
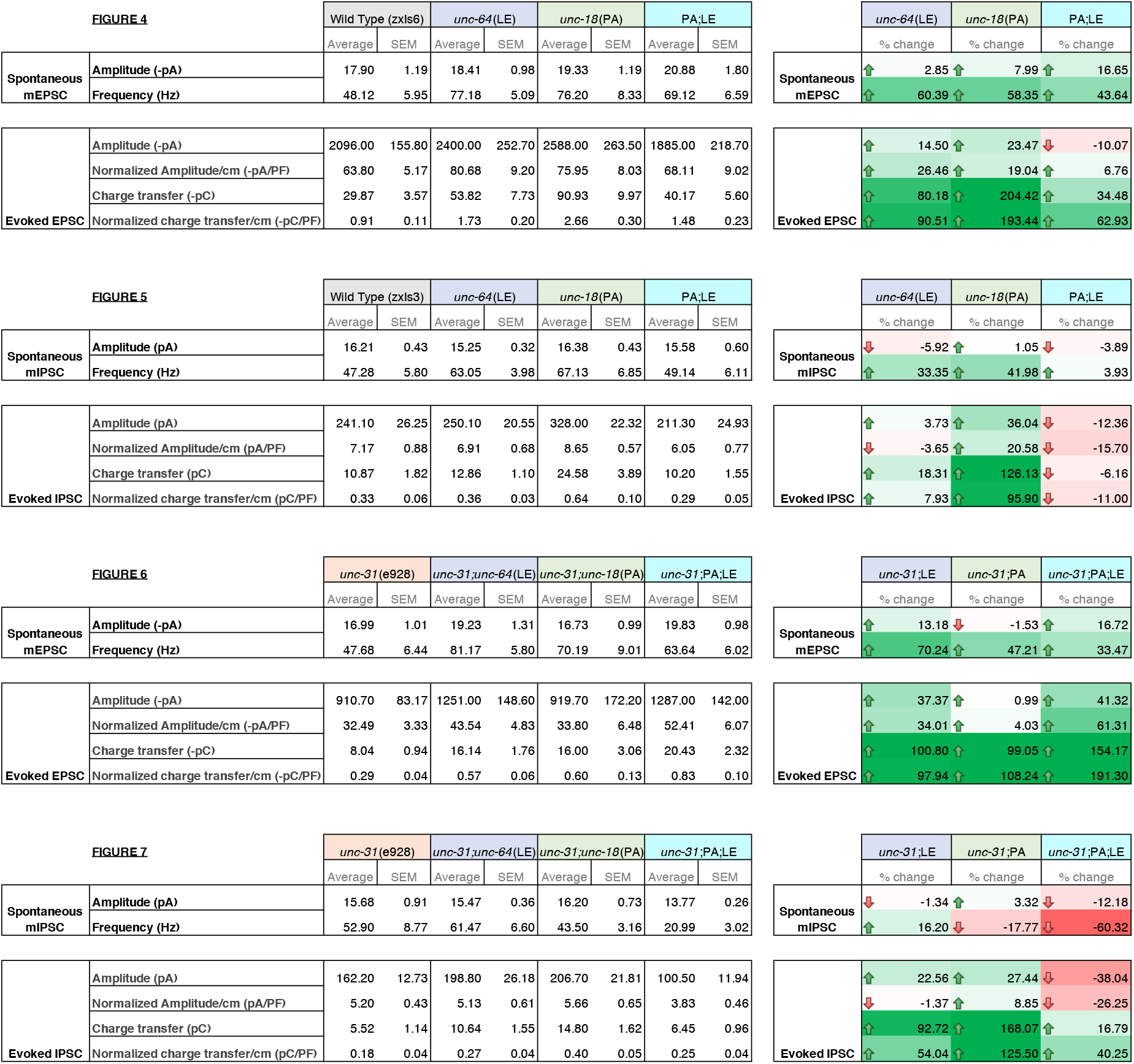
Summary of electrophysiology properties of indicated mutants. Summary values of electrophysiology data from Figures 3-7. Average values and standard error of the mean are indicated. Percent change for each category are indicated on the right with color scaling indicative of degree of change; +100% green, 0 white, -100% red.

### P334A UNC-18 enhances inhibitory synaptic transmission while open syntaxin and the double mutant do not

We then investigated inhibitory synaptic exocytosis using the single and double mutant strains in a *zxIs3* background. We found that, in the absence of light stimulation, the frequency of spontaneous miniature inhibitory postsynaptic currents (mIPSCs) was slightly increased in *unc-64(LE)* and *unc-18(PA)* worms (Fig. 5A, 5C). However, unlike mEPSCs, the frequency of mIPSCs of the *unc-64(LE); unc-18(PA)* double mutant did not show an increase (Fig. 5A, 5C, table 1). Again, amplitudes of mIPSCs were unchanged across all strains (Fig. 5B). We then looked at evoked inhibitory postsynaptic currents (IPSCs) by optogenetically stimulating the GABAergic neurons. We found that only the P334A UNC-18 mutant, *unc-18(PA),* facilitated IPSC charge transfer and amplitude, but this facilitation was lost in the *unc-64(LE); unc-18(PA)* double mutant (Fig. 5D-H). Not only was the facilitation lost, there was a tendency for decrease in averaged changes to both normalized and unnormalized amplitude and charge transfer (table 1). Thus, the double mutant fails to enhance spontaneous and evoked transmitter release in GABAergic inhibitory synapses, and may further emphasize inhibitory effects. Together with the data of excitatory transmitter release (Fig. 4), our results suggest that the double mutant of open syntaxin and P334A UNC-18 causes an imbalance between excitatory and inhibitory transmission. This may explain the phenotypes we observed where double mutants exhibited higher aldicarb sensitivity with reduced motility.

**Figure 5.**
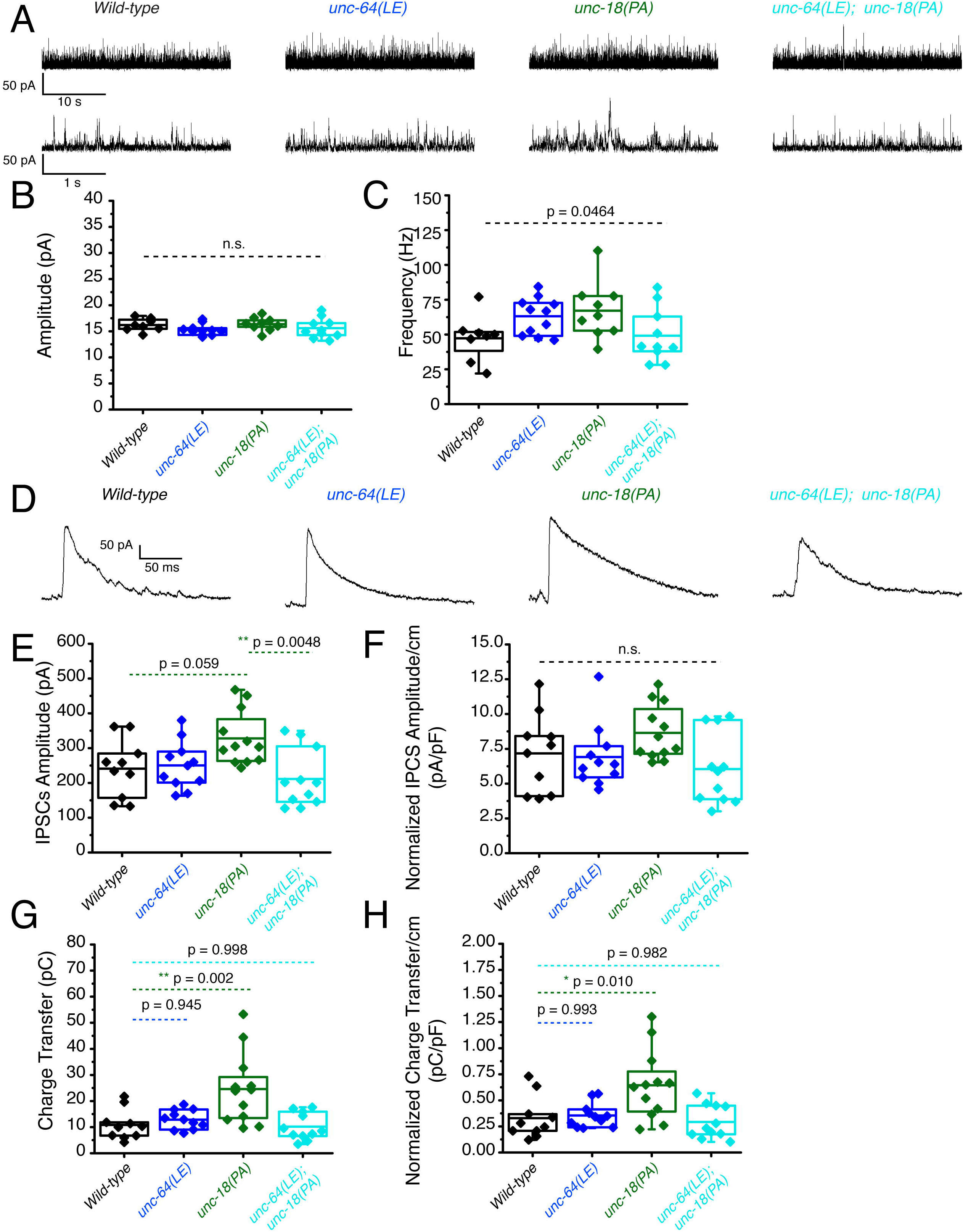
P334A unc-18 enhances inhibitory synaptic transmission while open syntaxin and the double mutant do not. (A) Sample traces of miniature inhibitory postsynaptic currents (mIPSCs) of wild-type (black), open syntaxin (unc-64(LE), blue), P334A UNC-18 (*unc-18(PA)*, green), and the resulting double mutant (cyan). (B) Amplitude of mIPSCs was unchanged across all strains (one-way ANOVA, F_(3, 34)_ = 1.369, p = 0.2687). (C) Frequency of mIPSCs. Both unc-18(PA) and unc-64(LE) slightly increase the frequency of mIPSCs, while the double mutant does not increase the frequency when compared to the wild-type (one-way ANOVA, F_(3, 34)_ = 2.952; p = 0.0464). (D) Sample traces of evoked inhibitory postsynaptic currents (IPSCs) for the indicated strains. (E) IPSC amplitude was increased by the unc-18(PA) mutant. However, this increase was lost in the unc-64(LE);unc-18(PA) double mutant (one-way ANOVA, F_(3, 40)_ = 4.742; p = 0.0064) (F) IPSC amplitude when calibrated with membrane capacitance to account for differences in animal size (one-way ANOVA, F_(3, 40)_ = 2.367; p = 0.0852). (G) Charge transfer of IPSCs. Only the unc-18(PA) single mutant showed an increase in charge transfer (one-way ANOVA, F_(3, 40)_ = 7.793; p = 0.0003); however, both unc-18(LE) or the unc-64(PA);unc-18(LE) double mutant did not show a significant increase in charge transfer. (H) Charge transfer of EPSCs calibrated to membrane capacitance to account for differences in animal size. Charge transfer again is increased for the unc-18(PA) single mutants. This increase was again lost in the unc-64(LE);unc-18(PA) double mutant (one-way ANOVA, F_(3, 40)_ = 6.132; p = 0.0016).

### P334A UNC-18, open syntaxin, and their double mutant all rescue reduced excitatory synaptic transmission in *unc-31* null worms

We showed that *unc-64(LE)* and *unc-18(PA)* can rescue the motility and aldicarb sensitivity defects of *unc-31(e928)* null mutants (Fig. 3A, 3B). However, the double mutant of *unc-64(LE); unc-18(PA)* worsened the mobility of *unc-31* mutants while restoring aldicarb to wild-type level (Fig. 3). Therefore, we analyzed these mutants in a *zxIs6* or *zxIs3* background using electrophysiology. *unc-31(e928)* null mutants did not alter mEPSC frequency compared with wild type worms (Fig. 6A, 6C). When the single mutants or the double *unc-64(LE); unc-18(PA)* mutant were crossed into the *unc-31(e928)* background, only *unc-31; unc64(LE)* significantly increased mEPSCs frequency (Fig. 6A, 6C). While trends for increased frequency were observed in the *unc-31; unc-18(PA)* and *unc-31; unc-64(LE); unc-18(PA)* worms, they did not reach statistical significance. Again, mEPSC amplitudes were unchanged (Fig. 6B). We then tested evoked release by optogenetic stimulation. *unc-31(e928)* exhibited decreased evoked EPSC amplitude and charge transfer compared with wild-type worms (Fig. 6D, 6E, 6G), which supports our recent findings that CAPS1/UNC-31 protein plays a role in synaptic vesicle release in addition to dense-core vesicle release (Wang, 2023; manuscript in preparation). Only *unc-31(e928)*; *unc-64(LE)* and *unc-31(e928); unc-64(LE); unc-18 (PA)* showed slightly increased evoked EPSC amplitudes with respect to *unc-31(e928)*, but without reaching statistical significance (Fig. 6D-6F, table 1). Moreover, the low charge transfer of *unc-31* showed rescue by all mutants: open syntaxin, P334A UNC-18, and the double mutant (Fig. 6G, 6H, table 1). Thus, open syntaxin and P334A UNC-18 appeared to have a small level of additivity in the rescue of evoked release at the excitatory synapse of *unc-31*.

**Figure 6.**
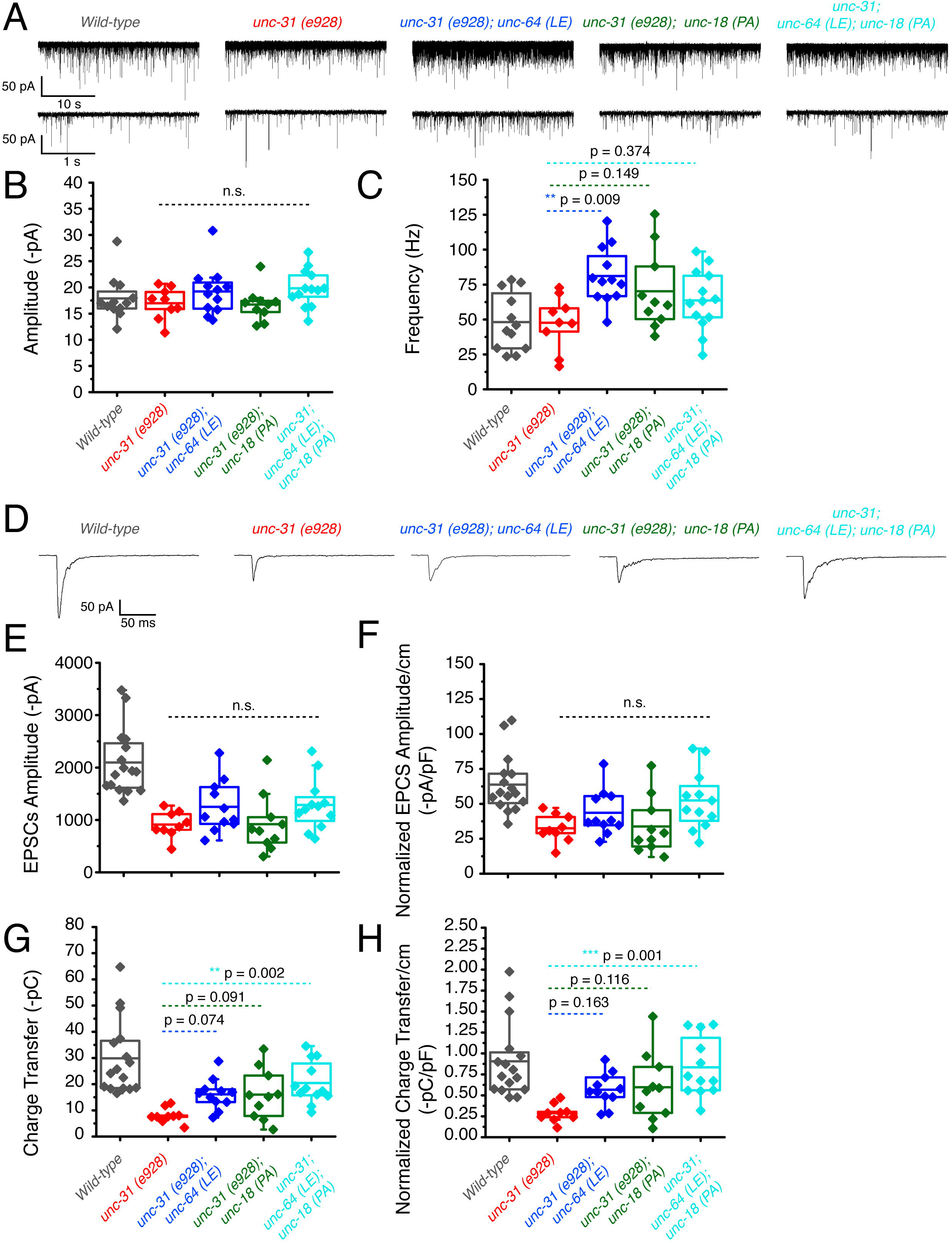
unc-31 null worms exhibit decreased EPSCs that are additively rescued by P334A unc-18 and open syntaxin. (A) Sample traces of spontaneous miniature postsynaptic currents (mPSCs) of wild-type (black), unc-31(e928), red, unc-31;*unc-64(LE)*, blue, unc-31;*unc-18(PA),*green and the resulting triple mutant (cyan). Wild-type data is from figure 4 for purpose of representation. (B) Amplitude of mPSCs was unchanged in all strains (one-way ANOVA, F_(3, 40)_ = 2.003; p = 0.1291). Wild-type data is from figure 4 for purpose of representation and excluded from analysis. (B) Frequency of mPSCs. Only *unc-31;unc-64(LE)* worms saw an increase in mPSC frequency (one-way ANOVA, F_(3, 40)_ = 3.933; p = 0.0150), while a trend of increase was observed in the *unc-31;unc-18(PA)* and *unc-31;unc-64(LE);unc-18(PA)* mutant. Wild-type data is from figure 4 for purpose of representation and excluded from analysis. (D) Sample traces of evoked excitatory postsynaptic currents (EPSCs) for the indicated strains. Wild-type data is from figure 4 for purpose of representation. (E) Evoked EPSC amplitude was reduced in the *unc-31* null animal when compared to control worms. A slight increase in EPSC amplitude was observed in the *unc-31*;*unc-64(LE)* and the *unc-31;unc-64(LE);unc-18(PA)* triple mutant but values did not reach statistical significance (one-way ANOVA, F_(3, 38)_ = 2.018; p = 0.1277). Wild-type data is from figure 4 for purpose of representation and excluded from analysis. (F) EPSC amplitude when calibrated with membrane capacitance to account for differences in animal size saw the same trend (one-way ANOVA, F_(3, 38)_ = 2.949; p = 0.0449). Wild-type data is from figure 4 for purpose of representation and excluded from analysis. (G) Charge transfer of EPSCs show a decrease in charge transfer of *unc-31* null worms when compared to wild-type worms. This decrease was rescued by the unc-31;unc-64(LE) and unc-31;unc-18(PA) double mutants, and further rescued by the unc-31;unc-64(LE);unc-18(PA) triple mutant (one-way ANOVA, F_(3, 38)_ = 5.178; p = 0.0042). Wild-type data is from figure 4 for purpose of representation and excluded from analysis. (H) Charge transfer of EPSCs calibrated to membrane capacitance to account for differences in animal size. Only *unc-31;unc-64(LE);unc-18(PA)* shows a significant rescue in EPSC charge transfer (one-way ANOVA, F_(3, 38)_ = 6.010; p = 0.0019). Wild-type data is from figure 4 for purpose of representation and excluded from analysis.

### P334A UNC-18 and open syntaxin rescue reduced inhibitory synaptic transmission of *unc-31* while the double mutant results in further impairment

Next, we investigated the inhibitory release potential of these worms by again crossing the various mutants into the *zxIs3* background. Spontaneous inhibitory release was unchanged in *unc-31(e928)* null worms as both frequency and amplitude were comparable to wild type levels (Fig. 7A-C). Neither *unc-31; unc-64(LE)* nor *unc-31; unc-18(PA)* double mutants largely changed these properties. However, there was a significant decrease in mIPSC frequency and amplitude in *unc-31; unc-64(LE); unc-18(PA)* triple mutants (Fig. 7A-7C, table 1). For evoked inhibitory release, *unc-31* null worms decreased in amplitude and charge transfer when compared to wild type controls (Fig. 7D, 7E, 7G). Both *unc-31; unc-64(LE)* and *unc-31; unc-18(PA)* worms showed trend of rescue in amplitude and charge transfer. By contrast, the *unc-31; unc-64(LE); unc-18(PA)* triple mutant exhibited a further decrease in amplitude when compared to *unc-31* null worms (Fig. 7D-7F, table 1). The rescuing effects on charge transfer that were observed in *unc-31; unc-64(LE)* and *unc-31; unc-18(PA)* worms were also lost in the triple mutant (Fig. 7G-7H, table 1). Compared with the data of excitatory acetylcholine transmitter release (Fig. 6), the evoked inhibitory data obtained with the *unc-31; unc-64(LE); unc-18(PA)* triple mutant suggest that there is an imbalance between excitatory and inhibitory transmitter release in both spontaneous and evoked release modes.

**Figure 7.**
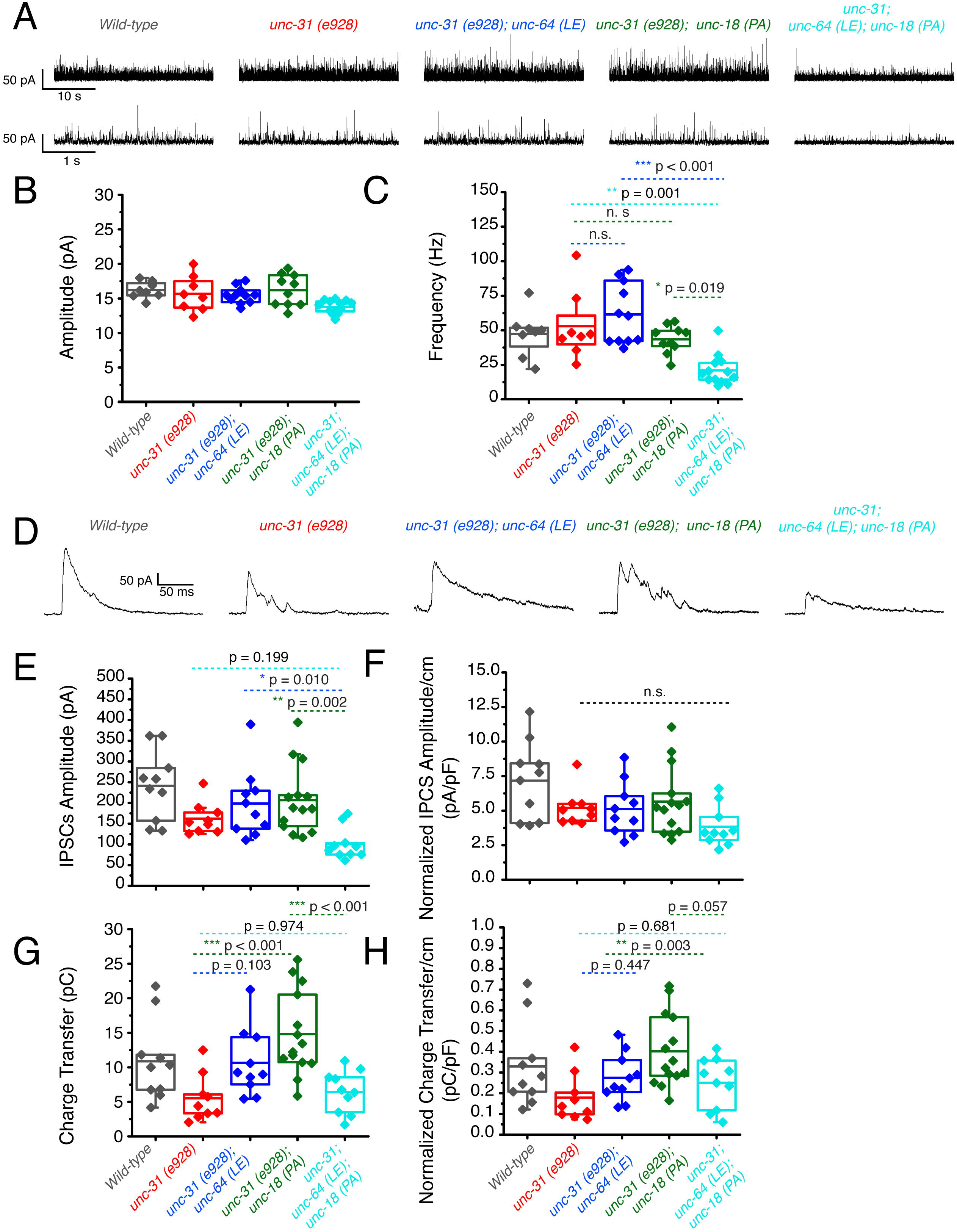
P334A unc-18 and open syntaxin rescues reduced inhibitory synaptic transmission of unc-31, but the double mutant does not. (A) Sample traces of miniature inhibitory postsynaptic currents (mIPSCs) of wild-type (black), *unc-31*(red), *unc-31;unc-64(LE) (*blue), *unc-31;unc-18(PA) (*green) and the resulting triple mutant (cyan). Wild-type data is from figure 5 for purpose of representation. (B) Amplitude of mIPSCs. The triple mutant showed a decrease in mIPSC amplitude (one-way ANOVA, F_(3, 38)_ = 4.163; p = 0.0121) that was significant from both the *unc-31;unc-64(LE)* and *unc-31*;*unc-18(PA)* double mutants. Wild-type data is from figure 5 for purpose of representation and excluded from analysis. (C) Frequency of mIPSCs. Both *unc-18(PA)* and *unc-64(LE)* do not alter the frequency of mIPSCS; however, the triple mutant decreased the frequency of mIPSCs (one-way ANOVA, F_(3, 38)_ = 12.04; p < 0.0001). Wild-type data is from figure 5 for purpose of representation and excluded from analysis. (D) Sample traces of evoked inhibitory postsynaptic currents (IPSCs) for the indicated strains. Wild-type data is from figure 5 for purpose of representation. (E) IPSC amplitude was decreased in *unc-31* null worms, while neither *unc-31;unc-64(LE)* or *unc-31;unc-18(PA)* significantly alleviated this deficit. The unc-31;unc-64(LE);unc-18(PA) triple mutant further reduced the size of IPSC amplitude (one-way ANOVA, F_(3, 39)_ = 5.713 ; p = 0.0024). Wild-type data is from figure 5 for purpose of representation and excluded from analysis. (F) IPSC amplitude when calibrated with membrane capacitance to account for differences in animal size (one-way ANOVA, F_(3, 39)_ = 1.831; p = 0.1575). Wild-type data is from figure 5 for purpose of representation and excluded from analysis. (G) Charge transfer of IPSCs found a decrease in charge transfer in the unc-31 null worms which was alleviated by both the *unc-31;unc-64(LE)* double mutant and the *unc-31;unc-18(PA)* double mutant. However, rescue effects were abolished in the *unc-31;unc-64(LE);unc-18(PA)* triple mutant (one-way ANOVA, F_(3, 39)_ = 9.456; p < 0.0001). Wild-type data is from figure 5 for purpose of representation and excluded from analysis. (H) Charge transfer of EPSCs calibrated to membrane capacitance to account for differences in animal size. Only *unc-31;unc-18(PA)* double mutant worms showed a significant rescue to charge transfer which was lost in the *unc-31;unc-64(LE);unc-18(PA)* triple mutant. One-way ANOVA, F_(3, 39)_ = 5.228; p = 0.0039. Wild-type data is from figure 5 for purpose of representation and excluded from analysis.

### The stimulatory effects caused by mammalian syntaxin-1 open and Munc18-1 P335A mutations on liposome fusion cancel each other

We found that the double mutation of open syntaxin and P334A UNC-18 in *C. elegans* has complex effects on exocytosis that depend on the type of (excitatory vs. inhibitory) synapse and on the mode of exocytosis (spontaneous vs. evoked) (Table 1). In short, the double mutant lost the ability to enhance inhibitory exocytosis displayed by the individual mutations. To investigate how combining the two mutations affects the ability of the neurotransmitter release machinery to induce membrane fusion in vitro, we used a well-defined liposome fusion assay that monitors lipid and content mixing, and uses mammalian proteins, including: i) the SNAREs syntaxin-1, SNAP-25 and synaptobrevin; ii) NSF and αSNAP, which disassemble SNARE complexes; and iii) Munc18-1 and a Munc13-1 C-terminal fragment spanning the executive region of this protein (Munc13-1C), which orchestrate SNARE complex assembly in a NSF-αSNAP-resistant manner (Ma et al., 2013; Liu et al., 2016; Stepien et al., 2019). This assay does not incorporate other regulator proteins such as tomosyn, complexin or CAPS, but recapitulates multiple central aspects of neurotransmitter release (Liu et al., 2016; Quade et al., 2019; Stepien and Rizo, 2021) as well as the gain-of-function effects of the syntaxin LE and P334A UNC-18 mutations (Park et al., 2017; Tien et al., 2020). These stimulatory effects are manifested in strong enhancements of Ca^2+^-independent fusion between liposomes containing syntaxin-1 (S-liposomes) and liposomes containing synaptobrevin (V-liposomes), which is barely observable when using WT proteins, whereas the high efficiency of fusion observed upon addition of Ca^2+^ does not allow the observation of stimulatory effects.

Indeed, control experiments with WT proteins revealed very slow fusion between S- and V-liposomes that was dramatically enhanced by Ca^2+^, whereas using P335A Munc18-1 led to a strong increase in Ca^2+^-independent fusion (Supplementary Fig. 1), as observed previously (Park et al., 2017). When we used S-liposomes containing open syntaxin-1 mutant, we observed a substantial amount of Ca^2+^-independent liposome fusion (Fig. 8) that reflects the gain-of-function caused by this mutation, as described previously (Tien et al., 2020). However, almost no Ca^2+^-independent liposome fusion was observed when we used P335A Munc18-1 and open syntaxin S-liposomes (Fig. 8), showing the stimulatory effects caused by the two mutations individually cancel each other. To shed light on into the molecular mechanism underlying these findings, we also performed liposome fusion assays with Munc18-1 bearing another mutation (D326K) that causes a gain-of-function because it helps to unfurl a Munc18-1 loop that covers the synaptobrevin binding site and, correspondingly, enhances synaptobrevin binding to Munc18-1 (Sitarska et al., 2017). D326K Munc18-1 enhanced liposome fusion in experiments performed with WT syntaxin-1 (Supplementary Fig. 1), as expected, and also enhanced Ca^2+^-independent fusion of open syntaxin-1 S-liposomes, in contrast to the P335A mutant (Fig. 8).

**Figure 8.**
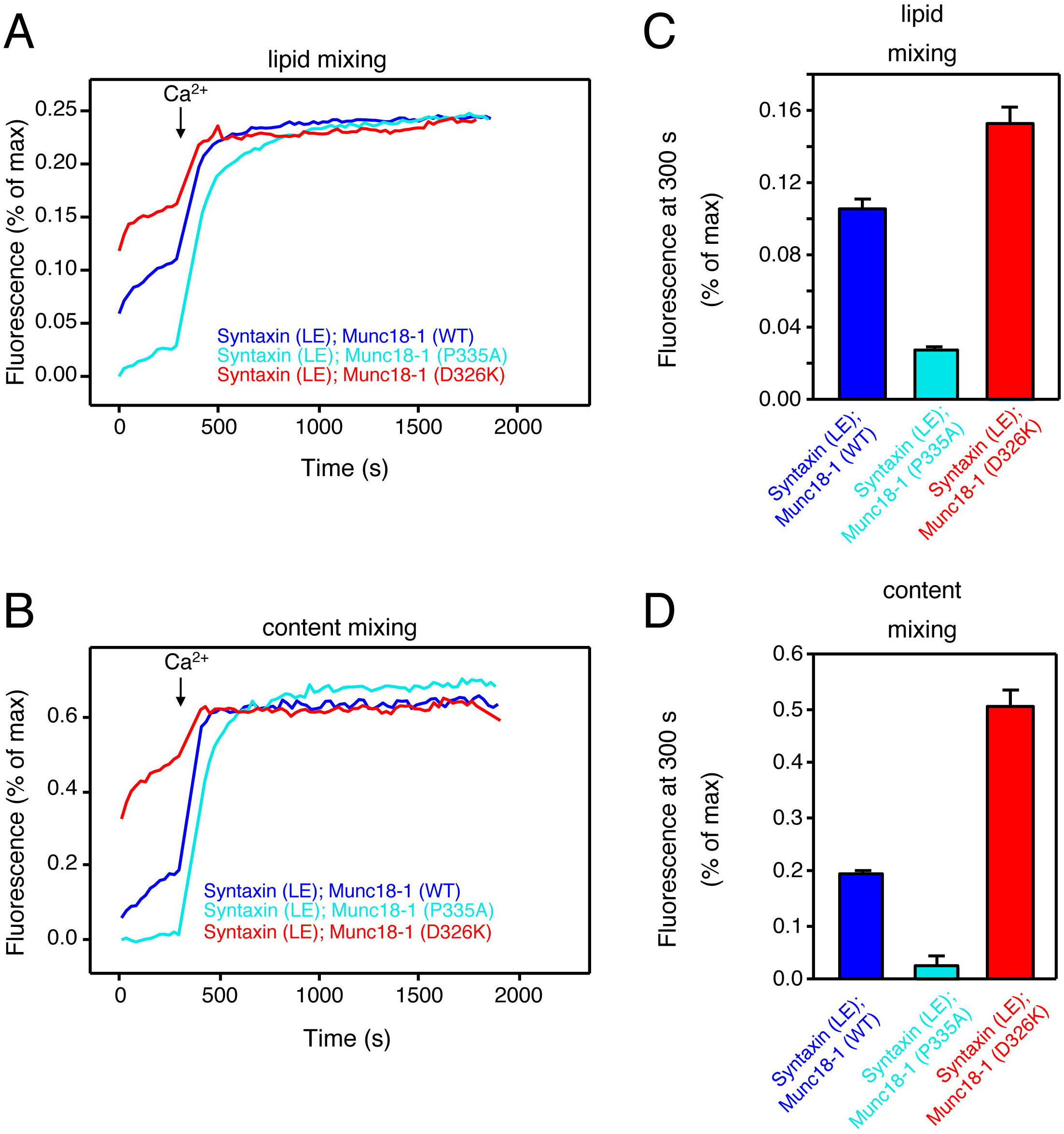
The stimulatory effects of the individual open syntaxin-1 and P335A Munc18-1 mutations on Ca^2+^-independent liposome fusion cancel each other in the double mutant. (A, B) Lipid mixing (A) between V-liposomes and S-liposomes containing open syntaxin-1 mutant was monitored from the fluorescence de-quenching of Marina Blue lipids, and content mixing (B) was monitored from the increase in the fluorescence signal of Cy5-streptavidin trapped in the V-liposomes caused by FRET with PhycoE-biotin trapped in the S-liposomes upon liposome fusion. Assays were performed in the presence of NSF, αSNAP, Munc13-1C and WT, P335A or D326K Munc18-1 as indicated by the color code. Experiments were started in the presence of 100 mM EGTA and 5 mM streptavidin, and Ca^2+^ (600 mM) was added at 300s. (C, D) Quantification of the fusion assays shown in panels A, B. Bars represent averages of the normalized fluorescence intensities observed for lipid mixing (C) or content mixing (D) at 300 s, performed in triplicates. Error bars represent standard deviations.

It is important to note that a key difference between the D326K and P335A Munc18-1 mutants is that the former does not affect syntaxin-1 binding but the latter substantially decreases the affinity of Munc18-1 for syntaxin-1 (Han et al., 2014; Stepien et al., 2022), and that the LE mutation that opens syntaxin-1 also causes a decrease in affinity for Munc18-1 (Dulubova et al., 1999; Stepien et al., 2022). We attempted to measure the affinity of P335A Munc18-1 for open syntaxin-1 using isothermal titration calorimetry, but the heats caused by binding were too small to yield accurate affinity measurements. Nevertheless, it is most likely that combining the two mutations further decreases the Munc18-1-syntaxin-1 affinity. Since αSNAP strongly inhibits liposome fusion and competes with Munc18-1 for binding to syntaxin-1 (Ma et al., 2013; Stepien et al., 2019), combining the open syntaxin-1 and P335A mutations is expected to tilt this competition in favor of αSNAP, leading to the cancellation of the stimulatory effects on liposome fusion caused by the individual mutations. In contrast, D326K Munc18-1 can still bind robustly to open syntaxin-1 and hence can further enhance the stimulatory effect of open syntaxin-1 on liposome fusion.

Overall, these observations show that the cancelation of the gains-of-function caused by the open syntaxin and P334A UNC-18 mutations in *C. elegans* can be recapitulated at least in part in liposome fusion assays performed with the mammalian proteins, and suggest that the differences in the physiological effects of the double mutation in excitatory and inhibitory synapses may arise because of differences in the relative levels of these various proteins (see below).

## Discussion

We found that the double mutation of open syntaxin and P334A UNC-18 has complex effects on exocytosis depending on the type of (excitatory vs. inhibitory) synapse and on the type (spontaneous vs. evoked) of exocytosis (Table 1). We first showed that P334A UNC-18 rescues detrimental phenotypes of the null *snt-1*(md290) mutant, which involves the major Ca^2+^ sensor protein for exocytosis (Fig. 1), similar to what we observed in open syntaxin mutants (Tien et al., 2020). We also found that similar to open syntaxin, P334A UNC-18 can also enhance exocytosis in a wide range of genetic backgrounds. In addition to rescuing *unc-13* null mutants (Park et al., 2017), P334A UNC-18 also rescues *snt-1* and *unc-31* mutants. These findings imply that P334A may provide a general means to enhance synaptic transmission in normal and disease states, much as the open syntaxin mutant does. Importantly however, the removal of tomosyn/TOM-1 did not rescue the exocytosis-deficient phenotype of *snt-1* null worms (Fig. 1). Tomosyn is considered to inhibit exocytosis by forming the inhibitory tomosyn-SNARE complex, and *tom-1* null worms enhance exocytosis by suppressing the formation of this inhibitory SNARE complex. On the other hand, open syntaxin and P334A UNC-18 enhance exocytosis by facilitating active SNARE assembly. Therefore, our results suggest that facilitation of the active SNARE assembly, but not the removal of inhibitory SNARE assembly, is needed to bypass the lack of synaptotagmin. This may imply that synaptotagmin is involved in active SNARE assembly, in agreement with results from in vitro assays of trans-SNARE complex formation (Prinslow et al., 2019).

Motivated by the rescue of a wide range of exocytosis-defective mutants by open syntaxin and P334A UNC-18 individually, we investigated the potential additional beneficial effects that could be caused by introducing both mutations simultaneously. However, we found that regardless of the genetic background, the double mutant had detrimental effects on many characteristics of the *C. elegans* that we tested, despite exhibiting an increased sensitivity to the acetylcholinesterase inhibitor, aldicarb (Fig. 1 - 3). Although the coincident occurrence of open syntaxin and P334A UNC-18 increased acetylcholine release, worms harboring both mutations did not have benefits for motility, body size, brood size, or growth speed.

Historically, in forward genetic screens, a large number of exocytosis defective mutants were isolated due to their resistance to cholinesterase inhibitors, which include aldicarb (Brenner, 1974; Nguyen et al., 1995; Miller et al., 1996). Therefore, aldicarb resistance and movement defects, such as the uncoordinated phenotype, are usually correlated in exocytosis mutants. In this context, our finding of the dissociation between motility and aldicarb sensitivity of the double mutant was surprising. This dissociation is particularly evident in the *unc-31* null background, as the animals bearing the open syntaxin and P334A UNC-18 mutations in this background exhibit aldicarb sensitivity similar to wild-type N2 levels, yet their thrashing is severely impaired (Fig. 3). These observations led us to study both excitatory and inhibitory synaptic transmission of the respective mutants in detail using electrophysiology combined with optogenetics (Fig. 4-7). As far as we know, this kind of detailed electrophysiological investigation in both spontaneous and evoked release in *C. elegans* excitatory and inhibitory synapses using optogenetics is unprecedented.

We summarized the synaptic transmission phenotypes of open syntaxin, P334A UNC-18 and the double mutant in the wild-type background and in the *unc-31* null background in Table 1. P334A UNC-18 consistently increased responses in excitatory and inhibitory evoked release in the wild-type and *unc-31* null background. Open syntaxin facilitated excitatory spontaneous and evoked release in both backgrounds, however, increases to inhibitory spontaneous and evoked transmission were weaker. The double mutant mirrored P334A UNC-18 in excitatory transmission and enhanced excitatory evoked charge transfer in both backgrounds. Strikingly however, the double mutant reduced spontaneous and evoked inhibitory release in both backgrounds. The differential effects of the double mutation on excitatory versus inhibitory synaptic transmission seem to explain why the double mutation worsened the motility of *unc-31* while rescuing aldicarb sensitivity. Thus, the imbalance of excitatory over inhibitory synaptic transmission increases acetylcholine release at the *C. elegans* neuromuscular junctions and increases aldicarb sensitivity, while impairing motility and other *C. elegans* features such as motility, brood size, or body size (Fig. 9). Our results also highlight the importance of investigating excitatory and inhibitory transmission when phenotypically examining exocytosis mutants in the future.

**Figure 9.**
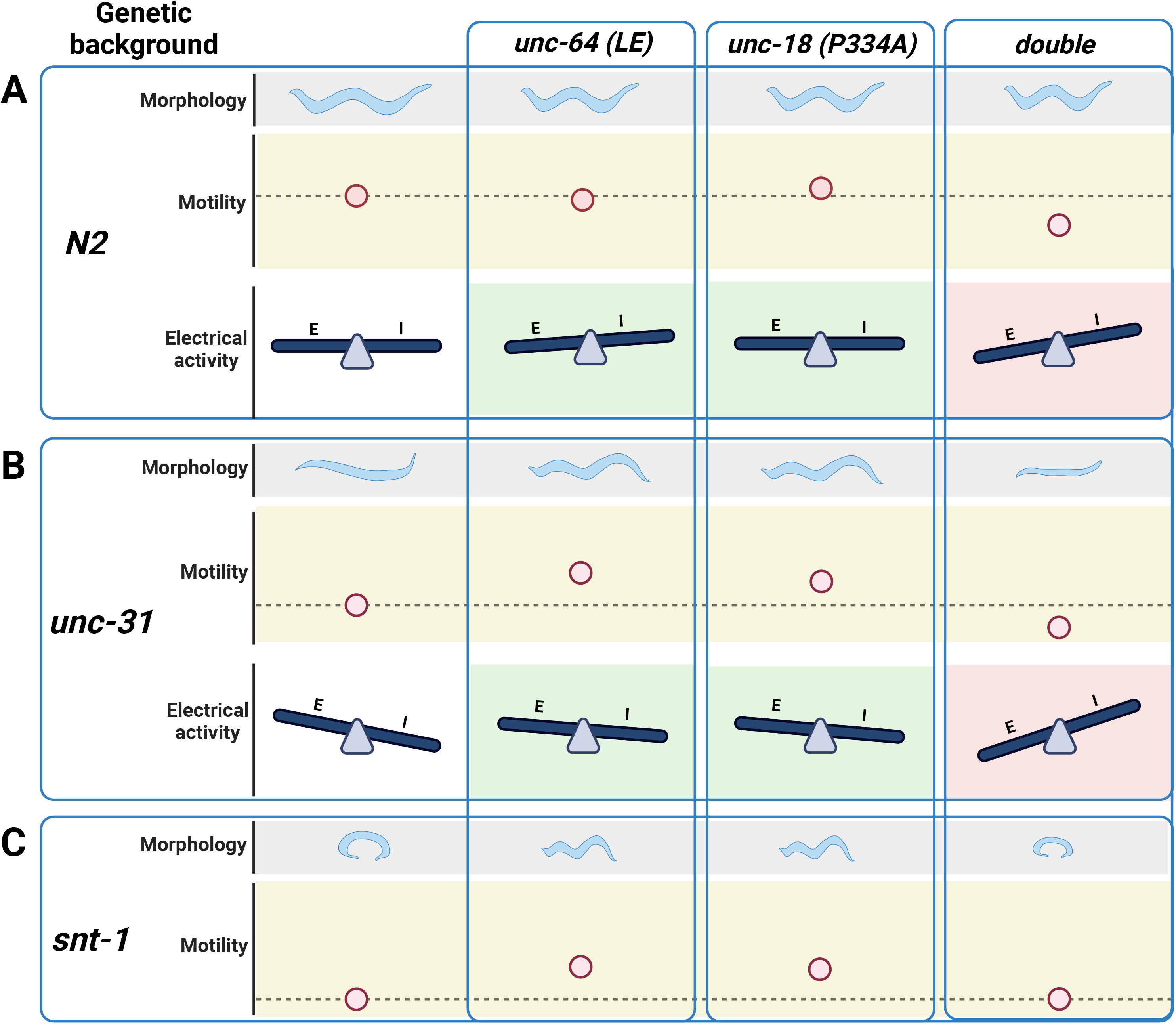
Summary of findings in open syntaxin, P334A UNC-18, and the simultaneous double mutant. (A) *C. elegans* characteristics such as worm size (morphology), and sinusoidal movement (motility) require proper coordination and balance of excitatory (E) and inhibitory (I) synaptic activity (electrical activity). While open syntaxin and P334A UNC-18 single mutations increase the excitatory behavior in *C. elegans* (green background), simultaneously open syntaxin and P334A UNC-18 mutations (double) decrease the size of the worm and the worm motility, and upsets the excitatory-inhibitory balance in a negative direction (red background). (B) *unc-31 null* worms exhibit a decrease in body size and motility, but these are rescued by open syntaxin and P334A UNC-18. Accordingly, both spontaneous and excitatory transmission are increased in the double mutants (green background). Simultaneous open syntaxin and P334A UNC-18 once again worsened worm features and motility, and disrupted the E-I balance in these worms (red background) (C) *snt-1 null* worms have a small body size and decreased motility. These are again rescued by both open syntaxin and P334A UNC-18; however, simultaneous presence of these mutants impaired this rescue. Figure created with BioRender.com

Our reconstitution results (Fig. 8), together with previous studies (Dulubova et al., 1999; Han et al., 2014; Park et al., 2017; Sitarska et al., 2017; Stepien et al., 2019; Tien et al., 2020; Stepien et al., 2022) suggests a model to explain the distinct phenotypes caused by the double open syntaxin, P334A UNC-18 mutation whereby neurotransmitter release depends on a delicate balance between the inhibitory activity of SNAPs and the crucial stimulatory functions of the SNARE complex assembly machinery formed by UNC-18 and UNC-13. Thus, studies of the mammalian proteins have shown that Munc18-1 and αSNAP compete for binding to syntaxin-1, and that binding of Munc18-1 to syntaxin-1 provides the only means to overcome the inhibition by αSNAP (Ma et al., 2013; Stepien et al., 2019). Since the affinity of the Munc18-1-syntaxin-1 complex is very high (in the low nanomolar range, (Burkhardt et al., 2008; Stepien et al., 2022)), the moderate impairments of binding caused by the open syntaxin-1 and P335A Munc18-1 mutations individually still allow strong binding (Burkhardt et al., 2008; Stepien et al., 2022) that readily overcomes the αSNAP inhibition (Park et al., 2017; Tien et al., 2020). In fact, these decreased affinities underlie the gains-of-function caused by these mutations, one because it helps to open syntaxin-1 (Dulubova et al., 1999) and the other because it facilitates formation of the synaptobrevin-Munc18-1-syntaxin-1 template complex that initiates SNARE complex assembly (Stepien et al., 2022). Note that trans-SNARE complex assembly with the WT proteins is inefficient in the absence of Ca^2+^ and is dramatically stimulated by Ca^2+^ (Prinslow et al., 2019), which underlies the strong Ca^2+^ dependence of liposome fusion in our reconstitution assays (Ma et al., 2013; Liu et al., 2016) (Supplementary Fig. 1). Ca^2+^-independent liposome fusion does occur more efficiently in experiments performed with open syntaxin-1 or P335A Munc18-1 because these mutations facilitate trans-SNARE complex assembly, but combining both mutations cancels these stimulatory effects of the individual mutations (Fig. 8, Supplementary Fig. 1), most likely because the combined decrease in affinity of the Munc18-1 for syntaxin-1 caused by both mutations tilts the balance in favor of αSNAP in its tug-of-war with Munc18-1. Nevertheless, there is still sufficient affinity between open syntaxin-1 and P335A Munc18-1 to form a functional complex, as liposome fusion still occurs in the presence of Ca^2+^, which dramatically enhances the efficiency of trans-SNARE complex assembly (Prinslow et al., 2019).

This model suggests that in *C. elegans* synapses the effects of the double mutation depends on the levels of the various proteins. For instance, the model predicts that, if the UNC-18 to SNAP ratio is low, combining the open syntaxin and P334A UNC-18 mutations will impair neurotransmitter release because of the impaired ability to overcome the SNAP inhibition. Conversely, if the UNC-18 to SNAP ratio is higher, the effects of combining the open syntaxin and P334A UNC-18 mutations are expected to be less deleterious. Note also that the effects of the double mutation likely depends also on the levels of UNC-13 and SNT-1, which are believed to synergize with UNC-18 in mediating trans-SNARE complex assembly based on the results obtained with mammalian proteins (Prinslow et al., 2019), and perhaps other proteins involved in SNARE assembly.

Overall, our results illustrate the complexity of the neurotransmitter release machinery and the different modes of regulation of release, and emphasize the importance of examining genetic interactions by generating double, triple and quadruple mutants. In this aspect, *C. elegans* as a model organism has significant advantages over mammalian systems. In mammals, closely related isoforms (e.g., syntaxin-1A, 1B; Munc13-1, 2; Tomosyn-1, 2) are present for each protein, and the compensatory and/or redundant effects of these isoforms makes it very difficult to evaluate the function of each protein (Fujiwara et al., 2006; Kofuji et al., 2014; Mishima et al., 2014). Conversely, even single knockout of certain proteins (e.g., syntaxin-1B, Munc18-1, Munc13-1) can cause embryonic or perinatal death in mice, which makes it virtually impossible to generate and analyze double or triple knockouts in mice. Thus, we believe that the detailed analysis of various *C. elegans* mutants using behavioral approaches and electrophysiology will keep providing new insights regarding the mechanisms of synaptic transmission as well as their relationship to behavior and other attributes of *C. elegans* life.

## Acknowledgements

This work was supported by the Natural Sciences and Engineering Research Council of Canada (RGPIN 2020 07139 to S.S), the Canadian Institute of Health Research (CIHR PJT 165917 to S.S), the Major International (Regional) Joint Research Project (32020103007 to S.G.), the National Key Research and Development Program of China (2022YFA1206001 to S.G.), the National Natural Science Foundation of China (31871069 to S.G.) and NIH Research Project Award R35 NS097333 to J.R.). M.H. and C.H.C. are supported by the Vision Science Research Program, Canada Graduate Scholarship, and the Ontario Graduate Scholarship.

## Author Contributions

M.H., Y.W., CH.C, and K.S, J.X., K.I, P.A., performed experiments, analyzed data and wrote the manuscript. X.X., K.S., C-W.T., S.L., contributed to experiments. S.G., S.S., J.R conceived the experiments, supervised the project, and assisted in writing the manuscript. P.P.M. discussed experiments and contributed reagents.

## Methods

### Worm maintenance

Worms were cultured using standard techniques. All strains used in the study were maintained at 22 °C on 30 mm agar NGM plates and seeded with OP50 as a food source. All C. elegans strains used are listed in Table 2.

**Table 2.**
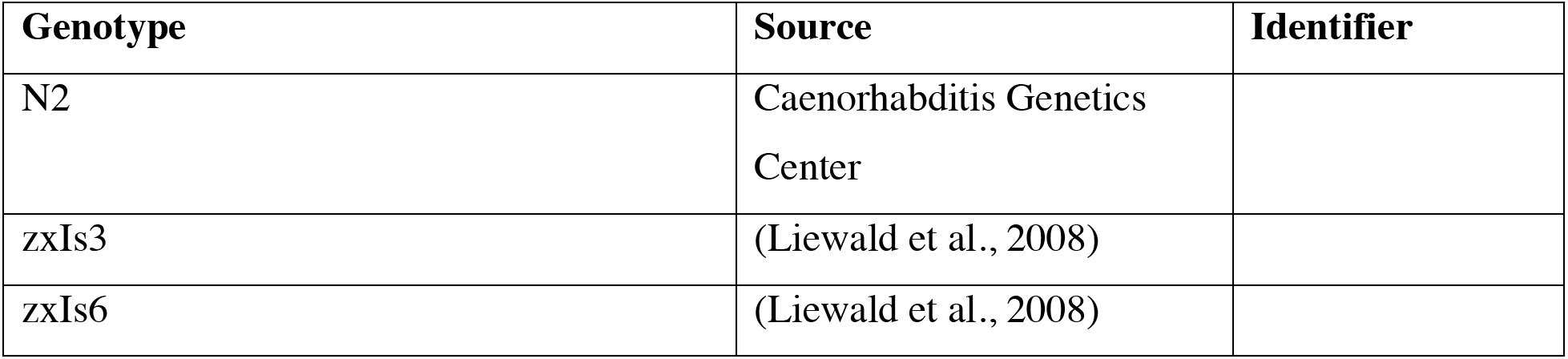
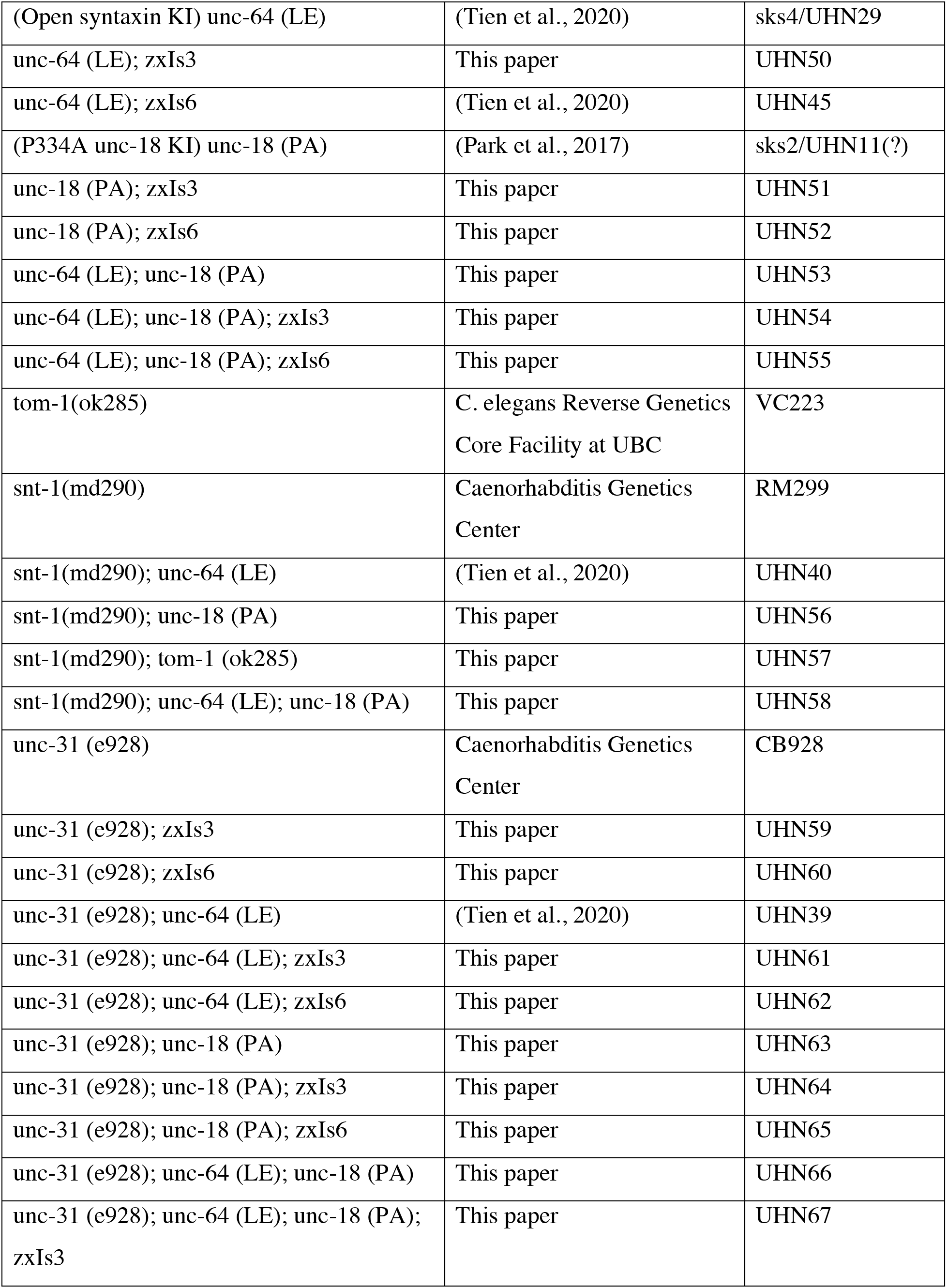
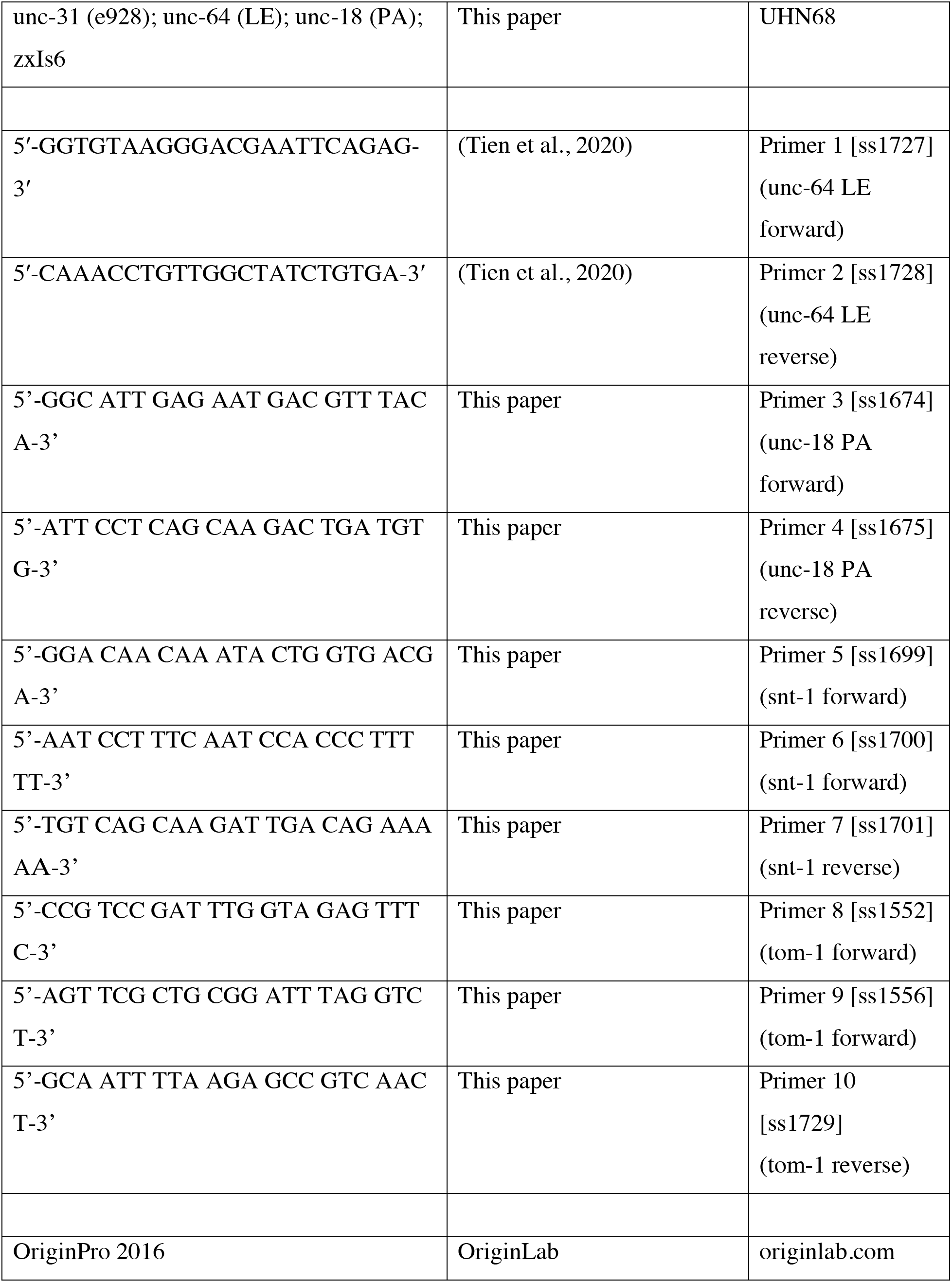
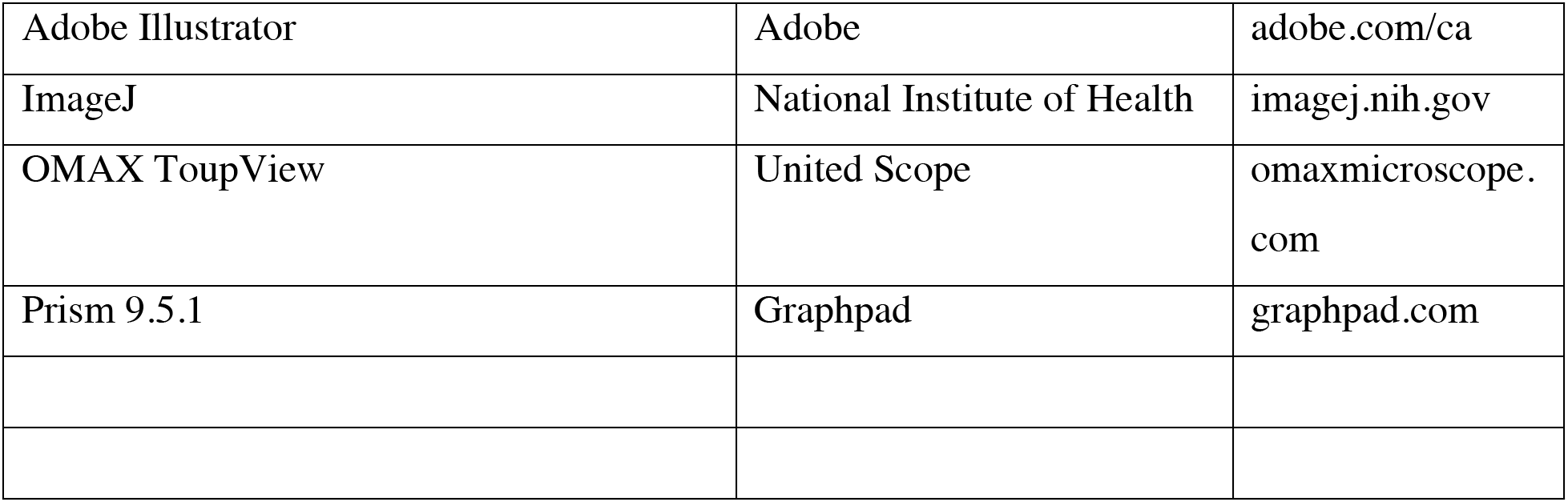
Worm strains and reagent used within this paper.

### Genetics

Fifteen to twenty adult worms from each plate were dissolved in 1X PCR buffer with 1mg/ml proteinase K to extract their DNA for PCR. PCR was conducted to confirm the genotype of double or triple mutants using primers purchased from IDT DNA. Primers are listed in the Table 2. A fluorescence scope was used to identify the zxIs3 and zxIs6 strains.

### Behavioural analyses

Worms were synchronized for all described assays. Briefly, gravid worms were bleached in a mixture of bleach and NaOH to release their eggs. Synchronized worms were allowed to grow to L4 stage/young adulthood before being assayed. In any assay spanning more than 4 hours, worms were grown within the same box on the same shelf during the course of the assay.

### Motility assays

Motility of each strain was determined by counting the thrashing rate of C. elegans in liquid medium. Post L4 young adult worms were placed in 60 μL of M9 buffer on a 35 mm petri dish lid. Worms were recorded for 4 minutes using an OMAX A3580U camera on a dissecting microscope with the OMAX ToupView v3.7 software. Worms were allowed to recover for 2 minutes and the later 2 minutes of each recorded video was used for thrashing analysis. Thrashes per minute was manually counted and averaged within each strain. A minimum of 40 worms per strain were used for analysis. A thrash was defined as a complete bend in the opposite direction at the midpoint of the body.

### Aldicarb assays

Aldicarb sensitivity was assessed using synchronously grown adult worms placed on non-seeded 35 mm NGM plates containing 0.3 mM or 1 mM aldicarb. All assays were done in 1 mM aldicarb plates unless specified otherwise. Over a 4 hour period, worms were monitored for paralysis at 15 or 30 min intervals, worms were checked a final time 24 hours later. Worms were considered paralyzed when there was no movement or pharyngeal pumping in response to three taps to the head and tail with a platinum wire. Once paralyzed, worms were removed from the plate. Six to eight sets of approximately 15 worms were examined for each strain.

### Worm size

Using synchronized worms, two days past the L4 stage, worms were placed in M9 liquid buffer with 50 mM sodium azide to prevent the worm from moving during image acquisition. The worm was imaged using a Nikon scope and then the perimeter of the worm was determined. The perimeter of the worm was manually traced using ImageJ. 20 worms were averaged for each strain.

### Population growth

From a synchronized population of worms, three young adult worms were moved to a new plate containing OP50. No worms with visibly present eggs were selected to ensure synchronicity. The number of L4 and above worms were counted daily until the whole plate reached starvation. Each strain was repeated a minimum of 6 times. All worms were grown in the same box on the same shelf through the duration of the experiment.

### Brood size

From a synchronized population of worms, a single L4 worm with a visible developing vulva but no eggs was isolated onto a new plate and let to lay eggs. 48 hours later the original worm was moved to a new plate. Following, the worm was moved to a new plate every 24 hours and this was repeated until egg laying ceased. Eggs and worms on the plate were counted after the mother worm was moved away. Two days after the original worm was moved away, unhatched eggs and worms on the plate were counted for the final count. Minimum replicate number was 6 for all strains, worms were grown in same box and same shelf throughout the duration of the experiment.

### Electrophysiology

The dissection and recording of the C. elegans was described previously (Gao and Zhen, 2011; Richmond and Jorgensen, 1999; Mellem et al., 2008). Briefly, 1 or 2 days old hermaphrodite adults were glued (Histoacryl Blue, Braun, Germany) to a sylgard (Sylgard 184, Dow Corning, USA)-coated cover glass covered with bath solution. The integrity of the ventral neuromuscular junction preparation was visually examined via DIC microscopy (Eclipse FN1, Nikon, Japan) after dissection, and muscle cells were patched using fire-polished 4–6 MΩ resistant borosilicate pipettes (1B100F-4, World Precision Instruments, USA). Membrane currents were recorded in the whole-cell configuration by PULSE software with a HEKA EPC-9 amplifier (Germany), and processed with Igor Pro 6.12 (WaveMetrics, USA) and Clampfit 11.0.3 (Axon Instruments, Molecular Devices, USA). Data were digitized at 10 kHz and filtered at 2.6 kHz.

Light stimulation of zxIs6 or zxIs3 was performed with a LED light source at a wavelength of 460 ± 5 nm (3.75 mW/mm2), triggered by the PULSE software with a duration of 10 ms. When recording mPSCs or eEPSCs, the muscle cells were held at -60 mV. To record mIPSCs or eIPSCs, 0.5 mM D-tubocurarine (d-TBC) were added to the bath solution to block all acetylcholine receptors with a holding potential of -10 mV. The recording solutions were described previously (Gao and Zhen). Specifically, the pipette solution contains (in mM): K-gluconate 115, KCl 25, CaCl_2_ 0.1, MgCl_2_ 5, BAPTA 1, HEPES 10, Na_2_ATP 5, Na_2_GTP 0.5, cAMP 0.5, cGMP 0.5, pH 7.2 with KOH, ∼320 mOsm. The bath solution consists of (in mM): NaCl 150, KCl 5, CaCl_2_ 5, MgCl_2_ 1, glucose 10, sucrose 5, HEPES 15, pH 7.3 with NaOH, ∼330 mOsm. All chemicals and blockers were from Sigma. Experiments were performed at room temperatures (20–22°C).

### Protein expression and purification

Bacterial expression and purification of full-length rat syntaxin-1A, the rat syntaxin L165E/E166A (LE) mutant (open syntaxin-1), a cysteine-free variant of full-length rat SNAP-25a, full-length rat synaptobrevin-2, full-length rat Munc18-1, full-length Chinese hamster NSF, full-length Bos Taurus αSNAP, and a fragments spanning the C-terminal region of rat Munc13-1 that includes the C_1_, C_2_B, MUN and C_2_C domains (residues 529-1725, Δ1408-1452, referred to as Munc13-1C) were described previously (Ma et al., 2013; Liu et al., 2016; Liu et al., 2017; Stepien et al., 2019). We note that WT or open syntaxin-1A were purified in buffer containing dodecylphosphocoline to prevent its aggregation (Liang et al., 2013).

### Lipid and content mixing assays

Assays that simultaneously measure lipid and content mixing between synaptobrevin-containing liposomes (V-liposomes) and syntaxin-1-containing liposomes (S-liposomes) were performed as previously described in detail (Stepien et al., 2019). Briefly, V-liposomes with reconstituted rat synaptobrevin-2 (protein-to-lipid ratio, 1:500) contained 40% POPC, 6.8% DOPS, 30.2% POPE, 20% cholesterol, 1.5% NBD PE, and 1.5% Marina Blue DHPE. S-liposomes with reconstituted WT or open rat syntaxin-1A (protein-to-lipid ratio, 1:800) contained 38% POPC, 18% DOPS, 20% POPE, 20% cholesterol, 2% PIP2, and 2% DAG. Lipid solutions were mixed with the respective proteins and with 4 μM Phycoerythrin-Biotin for S-liposomes or with 8 μM Cy5-Streptavidin for V-liposomes in 25 mM HEPES, pH 7.4, 150mM KCl, 1mM TCEP, 10% glycerol (v/v). To perform the lipid and content mixing assays, V-liposomes (0.125 mM lipids) were mixed with S-liposomes (0.25 mM lipids) in a total volume of 200 μL in the presence of 2.5 mM MgCl_2_, 2 mM ATP, 0.1 mM EGTA, 5 μM streptavidin, 0.4 μM NSF, 2 μM α-SNAP, 1 μM Munc18-1 (WT, P335A or D326K), 1 μM SNAP-25 and 0.5 μM Munc13-1C. Before mixing, S-liposomes were incubated with MgCl_2_, ATP, EGTA, streptavidin, NSF, αSNAP, and Munc18-1 at 37°C for 25 min. 0.6 mM of CaCl_2_ was added at 300 s to each reaction mixture. A PTI spectrofluorometer was used to measure lipid mixing from de-quenching of the fluorescence of Marina Blue–labeled lipids (excitation at 370 nm, emission at 465 nm) and content mixing from the development of FRET between PhycoE-Biotin trapped in the S-liposomes and Cy5-streptavidin trapped in the V-liposomes (PhycoE-biotin excitation at 565 nm, Cy5-streptavidin emission at 670 nm). All experiments were performed at 30°C. Lipid and content mixing were normalized as the percentage values of the maximum signals obtained by addition of 1% b-OG at the end to each reaction mixture (for lipid mixing) or to controls without streptavidin to measure maximal Cy5 fluorescence (for content mixing).

### Statistical analyses

All statistical analyses were done in OriginPro2016 and Prism 9.5.1. Independent t-tests were used for two-group experiments with a p-value < 0.05 as the threshold for statistical significance. One-way analysis of variance (ANOVA) was used for comparisons of multiple groups, followed by Tukey’s range test, with a significance level of 0.05. For all box and whisker plots, mean line is represented, range indicate 25 and 75 percentiles, whiskers are the outliers.

## References

Arunachalam L, Han L, Tassew NG, He Y, Wang L, Xie L, Fujita Y, Kwan E, Davletov B, Monnier PP, Gaisano HY, Sugita S (2008) Munc18-1 is critical for plasma membrane localization of syntaxin1 but not of SNAP-25 in PC12 cells. Mol Biol Cell 19:722–734.

Augustin I, Rosenmund C, Südhof TC, Brose N (1999) Munc13-1 is essential for fusion competence of glutamatergic synaptic vesicles. Nature 400:457–461.

Baker RW, Jeffrey PD, Zick M, Phillips BP, Wickner WT, Hughson FM (2015) A direct role for the Sec1/Munc18-family protein Vps33 as a template for SNARE assembly. Science 349:1111–1114.

Brenner S (1974) The genetics of Caenorhabditis elegans. Genetics 77:71–94.

Brose N, Hofmann K, Hata Y, Südhof TC (1995) Mammalian Homologues of Caenorhabditis elegans unc-13 Gene Define Novel Family of C2-domain Proteins (∗). Journal of Biological Chemistry 270:25273–25280.

Burkhardt P, Hattendorf DA, Weis WI, Fasshauer D (2008) Munc18a controls SNARE assembly through its interaction with the syntaxin N-peptide. Embo j 27:923–933.

Charlie NK, Schade MA, Thomure AM, Miller KG (2006) Presynaptic UNC-31 (CAPS) is required to activate the G alpha(s) pathway of the Caenorhabditis elegans synaptic signaling network. Genetics 172:943–961.

Dulubova I, Sugita S, Hill S, Hosaka M, Fernandez I, Südhof TC, Rizo J (1999) A conformational switch in syntaxin during exocytosis: role of munc18. The EMBO Journal 18:4372–4382.

Fernández-Chacón R, Königstorfer A, Gerber SH, García J, Matos MF, Stevens CF, Brose N, Rizo J, Rosenmund C, Südhof TC (2001) Synaptotagmin I functions as a calcium regulator of release probability. Nature 410:41–49.

Fujita Y, Shirataki H, Sakisaka T, Asakura T, Ohya T, Kotani H, Yokoyama S, Nishioka H, Matsuura Y, Mizoguchi A, Scheller RH, Takai Y (1998) Tomosyn: a Syntaxin-1–Binding Protein that Forms a Novel Complex in the Neurotransmitter Release Process. Neuron 20:905–915.

Fujiwara T, Mishima T, Kofuji T, Chiba T, Tanaka K, Yamamoto A, Akagawa K (2006) Analysis of knock-out mice to determine the role of HPC-1/syntaxin 1A in expressing synaptic plasticity. J Neurosci 26:5767–5776.

Geppert M, Goda Y, Hammer RE, Li C, Rosahl TW, Stevens CF, Südhof TC (1994) Synaptotagmin I: A major Ca2+ sensor for transmitter release at a central synapse. Cell 79:717–727.

Gerber SH, Rah JC, Min SW, Liu X, de Wit H, Dulubova I, Meyer AC, Rizo J, Arancillo M, Hammer RE, Verhage M, Rosenmund C, Südhof TC (2008) Conformational switch of syntaxin-1 controls synaptic vesicle fusion. Science 321:1507–1510.

Hammarlund M, Watanabe S, Schuske K, Jorgensen EM (2008) CAPS and syntaxin dock dense core vesicles to the plasma membrane in neurons. Journal of Cell Biology 180:483–491.

Han GA, Malintan NT, Saw NM, Li L, Han L, Meunier FA, Collins BM, Sugita S (2011) Munc18-1 domain-1 controls vesicle docking and secretion by interacting with syntaxin-1 and chaperoning it to the plasma membrane. Mol Biol Cell 22:4134–4149.

Han GA, Park S, Bin N-R, Jung CH, Kim B, Chandrasegaram P, Matsuda M, Riadi I, Han L, Sugita S (2014) A pivotal role for pro-335 in balancing the dual functions of Munc18-1 domain-3a in regulated exocytosis. The Journal of biological chemistry 289:33617–33628.

Han L, Jiang T, Han GA, Malintan NT, Xie L, Wang L, Tse FW, Gaisano HY, Collins BM, Meunier FA, Sugita S (2009) Rescue of Munc18-1 and -2 double knockdown reveals the essential functions of interaction between Munc18 and closed syntaxin in PC12 cells. Mol Biol Cell 20:4962–4975.

Hanson PI, Roth R, Morisaki H, Jahn R, Heuser JE (1997) Structure and conformational changes in NSF and its membrane receptor complexes visualized by quick-freeze/deep-etch electron microscopy. Cell 90:523–535.

Harrison SD, Broadie K, van de Goor J, Rubin GM (1994) Mutations in the Drosophila Rop gene suggest a function in general secretion and synaptic transmission. Neuron 13:555–566.

Hata Y, Slaughter CA, Südhof TC (1993) Synaptic vesicle fusion complex contains unc-18 homologue bound to syntaxin. Nature 366:347–351.

Hobson RJ, Liu Q, Watanabe S, Jorgensen EM (2011) Complexin maintains vesicles in the primed state in C. elegans. Curr Biol 21:106–113.

Hosono R, Hekimi S, Kamiya Y, Sassa T, Murakami S, Nishiwaki K, Miwa J, Taketo A, Kodaira KI (1992) The unc-18 gene encodes a novel protein affecting the kinetics of acetylcholine metabolism in the nematode Caenorhabditis elegans. J Neurochem 58:1517–1525.

Huntwork S, Littleton JT (2007) A complexin fusion clamp regulates spontaneous neurotransmitter release and synaptic growth. Nat Neurosci 10:1235–1237.

Jockusch WJ, Speidel D, Sigler A, Sorensen JB, Varoqueaux F, Rhee JS, Brose N (2007) CAPS-1 and CAPS-2 are essential synaptic vesicle priming proteins. Cell 131:796–808.

Jorgensen EM, Hartwieg E, Schuske K, Nonet ML, Jin Y, Horvitz HR (1995) Defective recycling of synaptic vesicles in synaptotagmin mutants of Caenorhabditis elegans. Nature 378:196–199.

Jospin M, Qi YB, Stawicki TM, Boulin T, Schuske KR, Horvitz HR, Bessereau J-L, Jorgensen EM, Jin Y (2009) A Neuronal Acetylcholine Receptor Regulates the Balance of Muscle Excitation and Inhibition in Caenorhabditis elegans. PLOS Biology 7:e1000265.

Kofuji T, Fujiwara T, Sanada M, Mishima T, Akagawa K (2014) HPC-1/syntaxin 1A and syntaxin 1B play distinct roles in neuronal survival. J Neurochem 130:514–525.

Koushika SP, Richmond JE, Hadwiger G, Weimer RM, Jorgensen EM, Nonet ML (2001) A post-docking role for active zone protein Rim. Nat Neurosci 4:997–1005.

Lee BH, Liu J, Wong D, Srinivasan S, Ashrafi K (2011) Hyperactive neuroendocrine secretion causes size, feeding, and metabolic defects of C. elegans Bardet-Biedl syndrome mutants. PLoS Biol 9:e1001219.

Li L, Liu H, Krout M, Richmond JE, Wang Y, Bai J, Weeratunga S, Collins BM, Ventimiglia D, Yu Y, Xia J, Tang J, Liu J, Hu Z (2021) A novel dual Ca2+ sensor system regulates Ca2+-dependent neurotransmitter release. Journal of Cell Biology 220.

Liang B, Kiessling V, Tamm LK (2013) Prefusion structure of syntaxin-1A suggests pathway for folding into neuronal trans-SNARE complex fusion intermediate. Proc Natl Acad Sci U S A 110:19384–19389.

Liewald JF, Brauner M, Stephens GJ, Bouhours M, Schultheis C, Zhen M, Gottschalk A (2008) Optogenetic analysis of synaptic function. Nat Methods 5:895–902.

Liu X, Seven AB, Xu J, Esser V, Su L, Ma C, Rizo J (2017) Simultaneous lipid and content mixing assays for in vitro reconstitution studies of synaptic vesicle fusion. Nat Protoc 12:2014–2028.

Liu X, Seven AB, Camacho M, Esser V, Xu J, Trimbuch T, Quade B, Su L, Ma C, Rosenmund C, Rizo J (2016) Functional synergy between the Munc13 C-terminal C1 and C2 domains. Elife 5.

Ma C, Li W, Xu Y, Rizo J (2011) Munc13 mediates the transition from the closed syntaxin-Munc18 complex to the SNARE complex. Nat Struct Mol Biol 18:542–549.

Ma C, Su L, Seven AB, Xu Y, Rizo J (2013) Reconstitution of the vital functions of Munc18 and Munc13 in neurotransmitter release. Science 339:421–425.

Martin JA, Hu Z, Fenz KM, Fernandez J, Dittman JS (2011) Complexin has opposite effects on two modes of synaptic vesicle fusion. Curr Biol 21:97–105.

McEwen JM, Madison JM, Dybbs M, Kaplan JM (2006) Antagonistic Regulation of Synaptic Vesicle Priming by Tomosyn and UNC-13. Neuron 51:303–315.

Miller KG, Alfonso A, Nguyen M, Crowell JA, Johnson CD, Rand JB (1996) A genetic selection for Caenorhabditis elegans synaptic transmission mutants. Proceedings of the National Academy of Sciences 93:12593–12598.

Mishima T, Fujiwara T, Sanada M, Kofuji T, Kanai-Azuma M, Akagawa K (2014) Syntaxin 1B, but not syntaxin 1A, is necessary for the regulation of synaptic vesicle exocytosis and of the readily releasable pool at central synapses. PLoS One 9:e90004.

Misura KMS, Scheller RH, Weis WI (2000) Three-dimensional structure of the neuronal-Sec1–syntaxin 1a complex. Nature 404:355–362.

Munch AS, Kedar GH, van Weering JR, Vazquez-Sanchez S, He E, Andre T, Braun T, Sollner TH, Verhage M, Sorensen JB (2016) Extension of Helix 12 in Munc18-1 Induces Vesicle Priming. J Neurosci 36:6881–6891.

Nguyen M, Alfonso A, Johnson CD, Rand JB (1995) Caenorhabditis elegans mutants resistant to inhibitors of acetylcholinesterase. Genetics 140:527–535.

Parisotto D, Pfau M, Scheutzow A, Wild K, Mayer MP, Malsam J, Sinning I, Söllner TH (2014) An Extended Helical Conformation in Domain 3a of Munc18-1 Provides a Template for SNARE (Soluble *N*-Ethylmaleimide-sensitive Factor Attachment Protein Receptor) Complex Assembly *. Journal of Biological Chemistry 289:9639–9650.

Park S et al. (2017) UNC-18 and Tomosyn Antagonistically Control Synaptic Vesicle Priming Downstream of UNC-13 in Caenorhabditis elegans.

Prinslow EA, Stepien KP, Pan YZ, Xu J, Rizo J (2019) Multiple factors maintain assembled trans-SNARE complexes in the presence of NSF and αSNAP. Elife 8.

Quade B, Camacho M, Zhao X, Orlando M, Trimbuch T, Xu J, Li W, Nicastro D, Rosenmund C, Rizo J (2019) Membrane bridging by Munc13-1 is crucial for neurotransmitter release. Elife 8.

Renden R, Berwin B, Davis W, Ann K, Chin CT, Kreber R, Ganetzky B, Martin TF, Broadie K (2001) Drosophila CAPS is an essential gene that regulates dense-core vesicle release and synaptic vesicle fusion. Neuron 31:421–437.

Richmond JE, Jorgensen EM (1999) One GABA and two acetylcholine receptors function at the C. elegans neuromuscular junction. Nat Neurosci 2:791–797.

Richmond JE, Davis WS, Jorgensen EM (1999) UNC-13 is required for synaptic vesicle fusion in C. elegans. Nat Neurosci 2:959–964.

Richmond JE, Weimer RM, Jorgensen EM (2001) An open form of syntaxin bypasses the requirement for UNC-13 in vesicle priming. Nature 412:338–341.

Rizo J (2022) Molecular Mechanisms Underlying Neurotransmitter Release. Annu Rev Biophys 51:377–408.

Sitarska E, Xu J, Park S, Liu X, Quade B, Stepien K, Sugita K, Brautigam CA, Sugita S, Rizo J (2017) Autoinhibition of Munc18-1 modulates synaptobrevin binding and helps to enable Munc13-dependent regulation of membrane fusion. Elife 6.

Stepien KP, Rizo J (2021) Synaptotagmin-1-, Munc18-1-, and Munc13-1-dependent liposome fusion with a few neuronal SNAREs. Proc Natl Acad Sci U S A 118.

Stepien KP, Prinslow EA, Rizo J (2019) Munc18-1 is crucial to overcome the inhibition of synaptic vesicle fusion by αSNAP. Nat Commun 10:4326.

Stepien KP, Xu J, Zhang X, Bai X-C, Rizo J (2022) SNARE assembly enlightened by cryo-EM structures of a synaptobrevin–Munc18-1–syntaxin-1 complex. Science Advances 8:eabo5272.

Sutton RB, Fasshauer D, Jahn R, Brunger AT (1998) Crystal structure of a SNARE complex involved in synaptic exocytosis at 2.4 Å resolution. Nature 395:347–353.

Tien C-W, Yu B, Huang M, Stepien KP, Sugita K, Xie X, Han L, Monnier PP, Zhen M, Rizo J, Gao S, Sugita S (2020) Open syntaxin overcomes exocytosis defects of diverse mutants in C. elegans. Nature Communications 11:1–18.

Verhage M, Maia AS, Plomp JJ, Brussaard AB, Heeroma JH, Vermeer H, Toonen RF, Hammer RE, den TKv, Berg, Missler M, Geuze HJ, Südhof TC (2000) Synaptic Assembly of the Brain in the Absence of Neurotransmitter Secretion.

Voets T, Toonen RF, Brian EC, de Wit H, Moser T, Rettig J, Sudhof TC, Neher E, Verhage M (2001) Munc18-1 promotes large dense-core vesicle docking. Neuron 31:581–591.

Weber T, Zemelman BV, McNew JA, Westermann B, Gmachl M, Parlati F, Söllner TH, Rothman JE (1998) SNAREpins: minimal machinery for membrane fusion. Cell 92:759–772.

Weimer RM, Richmond JE, Davis WS, Hadwiger G, Nonet ML, Jorgensen EM (2003) Defects in synaptic vesicle docking in unc-18 mutants. Nature Neuroscience 6:1023–1030.

