## Supplemental Files for "Double mutation of open syntaxin and UNC-18 P334A leads to excitatory-inhibitory imbalance and impairs multiple aspects of *C. elegans* behavior"

**SUPPLEMENTARY INFO**

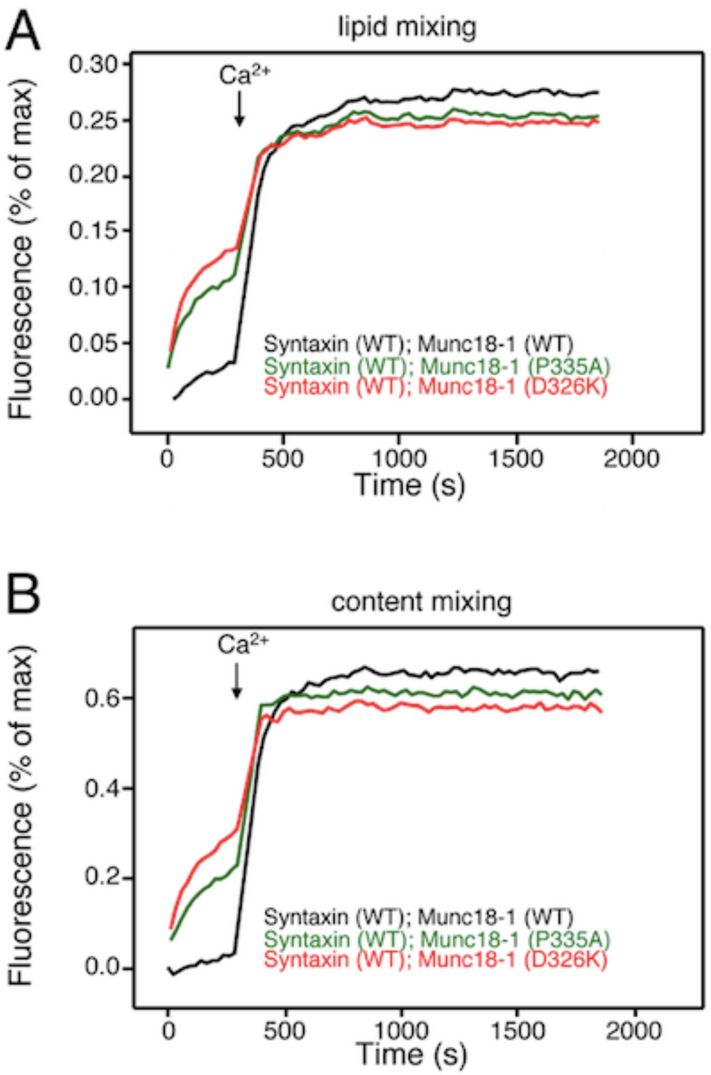


**Supplementary Figure 1.** Stimulatory effects of the P335A and D326K Munc18-1 mutations on $Ca^{2+}$-independent liposome fusion.

(A, B) Lipid mixing (A) between V-liposomes and S-liposomes containing WT syntaxin-1 was monitored from the fluorescence de-quenching of Marina Blue lipids, and content mixing (B) was monitored from the increase in the fluorescence signal of Cy5-streptavidin trapped in the V-liposomes caused by FRET with PhycoE-biotin trapped in the S-liposomes upon liposome fusion. Assays were performed in the presence of NSF, αSNAP, Munc13-1C and WT, P335A or D326K Munc18-1 as indicated by the color code. Experiments were started in the presence of 100 mM EGTA and 5 mM streptavidin, and $Ca^{2+}$ (600 mM) was added at 300s.
